# A microbial seedbank from a natural oil seep accelerates hydrocarbon degradation in freshwater oil spills

**DOI:** 10.64898/2026.03.16.712059

**Authors:** Rudolf Walter, Lea Willemsen, Lisa Voskuhl

## Abstract

Anthropogenic oil spills are a major source of hydrocarbon pollution, yet freshwater oil spills are studied less than marine once. We hypothesize that microorganisms from freshwater natural oil seeps are adapted to use oil as carbon source and could serve as a seedbank to accelerate degradation of anthropogenic oil spills.

To test this, microcosms were set up using water and oil from a natural oil seep in Germany, while light oil and oil naïve river water simulated an anthropogenic spill. Seven conditions combined different water types (adapted, naïve) and oil types (heavy, light, mixed). Oil degradation was monitored over 81 days via reverse stable isotope labeling, and pH and microbial communities were analyzed by 16S rRNA gene amplicon sequencing at the start and end. Microcosms with adapted microbes exhibited higher oil mineralization rates. In mixed-oil incubations, simulating the application of a microbial seedbank, degradation rates were 0.26 mM/day with adapted microbes versus 0.15 mM/day in naïve water. Applying seep-derived microbes to naïve light-oil microcosms increased hydrocarbon mineralization by 32%. These results suggest that leveraging natural microbial seedbanks could accelerate oil weathering and reduce the environmental impact of freshwater oil spills, offering a promising strategy for spill mitigation if applied safely.

**Importance:** Freshwater oil spills remain poorly studied despite their substantial contribution to global hydrocarbon pollution. This study demonstrates that microbial communities from freshwater natural oil seeps are pre-adapted to degrade crude oil and can significantly enhance the breakdown of anthropogenic oil spills. Applying such microbial seedbanks increased hydrocarbon mineralization by over 30%, highlighting their potential as an effective and environmentally relevant tool for oil spill mitigation. These findings support the development of nature-based strategies to accelerate oil weathering and reduce the environmental impact of freshwater oil spills.

## Introduction

Oil or hydrocarbons can enter the environment through different ways, either via natural seeps, found at tectonic plate boundaries or along cracks and faults in geological formations allowing the entrapped crude oil seeping from the underground to the surface (1). Or, oil enters via anthropogenic spills of crude or refined oil for example after shipwrecks or discharges, broken pipelines or leaks (2). When oil spills into a waterbody, the released oil spreads out and floats at the water surface and appears as an oil film on it, also called oil slick (3). In a study where synthetic aperture radar (SAR) images of global seas and oceans were analyzed, 435 natural slick forming oil seeps were identified. Natural leakages only contributed to 6.2% of the total oil slick area. As for anthropogenic oil input, pipelines (0.5%), platforms (1.6%), and other anthropogenic sources (e.g. ships, land-based discharges and other spills, 91.7%) attributed for a larger oil slick area (4). Such global numbers are to the best of our knowledge not available for freshwater systems, because here the application of satellite-based detection is less common but emerging (5, 6). Oil slicks persist and interact with the air-water interface, disturbing biological processes in this sensitive environment by altering light availability, nutrient transport, gas exchange, and the release of toxic compounds (3). Most research on oil slick microbiology focusses on marine systems and on anthropogenically caused spills, whereas freshwater environments such as lakes, rivers and streams and natural oil seeps receive comparatively little attention. This bias reflects most likely the perception that anthropogenic marine oil spills have greater global impact due to scale and frequency. However, oil pollution also occurs in freshwater systems where it can cause severe ecological and economic consequences but still remains insufficiently studied and thus requires more investigation. Examples of larger oil spill into fresh waters are the River Lambro’s (Italy) oil spill in 2010 with approximate 2.6 kilotons of oil spilled that also flowed into the River Po (7), chronic leaks from oil extractions into the Niger Delta, Nigeria (8) with an estimate of 1.5 million tons of oil spilled over a 50-year period (9), or the Kalamazoo River oil spill in Michigan, USA (2010) from a pipeline burst that released 3.1 million liters of oil into the environment (10). In addition, numerous small and often unreported oil releases occur in freshwater environments, for instance through river shipping, leakages in recreational marinas or runoff from roads following traffic accidents (11, 12). The severity of an oil spill most likely depends on the type of released oil, how long the ecosystems is exposed and how much oil was released (13). Harmful impacts could be mitigated if the oil becomes less toxic via rapid degradation or if it is degraded faster before reaching the shoreline, where it might otherwise also affect terrestrial organisms and vegetation (14). To protect the environment from the harmful impacts of oil spills, specialized oil spill response strategies have been implemented worldwide. Oil spill response aims to detect, contain, and mitigate oil spills to protect ecosystems, human health, and economies. The Exxon Valdez oil tanker spill was maybe the first full-scale and systematically evaluated field application of bioremediation where nutrient amendments were used to stimulate indigenous hydrocarbon-degrading bacteria (15). Approximately 41 million liters of crude oil were released during this spill. During the years 1989–1992, fertilizers (50 Mg of nitrogen and 5 Mg of phosphorus) was applied to enhance microbial biodegradation of the oil (16). Monitoring showed a mean loss in the mass of residual oil of about 28% per year for surface oil and 12% per year for sub-surface oil (15). Of the estimated existing 2.2–4.3 million prokaryotic OTUs (17) only a few hundred are known to be capable of hydrocarbon biodegradation (18), but also, they are known for being ubiquitous and naturally present in diverse environments (19). Natural microbial communities often harbor a diverse seedbank of dormant or low-abundant taxa that can become metabolically active and increase in numbers when environmental conditions shift and create more favorable environments for these organisms (20). This was shown for hydrocarbon degrading prokaryotes which have been found to occur in low abundances in unpolluted environments and rapidly, within hours to days, responded to oil exposure by increased growth and accelerated biodegradation (21–24). Based on the ecological seedbank concept, microbial communities that are adapted to chronic oil exposure in environments, such as natural oil seeps or slicks, could act as functional seedbanks, carrying metabolic capabilities for oil degradation. When such a seedbank is transferred to freshly contaminated sites, these consortia could potentially establish rapid oil degradation, effectively priming the recipient environment for biodegradation. This principle is analogous to microbial inoculation or bioaugmentation, but it emphasizes ecological compatibility and resilience derived from prior adaptation (25, 26). Using microbial seedbanks that have co-evolved with hydrocarbons for long times enables remediation strategies to exploit the genetic potential and cooperative interactions that have been optimized through natural selection.

The aim of this study was to test if **i)** microorganisms that were isolated from a natural oil seep into freshwater, are strongly adapted to hydrocarbon degradation and **ii)** can potentially be applied to anthropogenic oil spills into freshwater that was naïve to oil before, and to increase the natural oil degradation rates. In other words, if microorganisms from natural oil slicks can serve as seedbanks for faster biodegradation of anthropogenic oil spills into recently contaminated freshwaters. To test our hypotheses, we took environmental samples from a potentially adapted microbiome isolated from a natural oil seep in Germany, where heavy oil seeps into groundwater for more than 500 years and samples from the river Aller, that contain microorganisms naïve to chronic oil exposure. We performed microcosm experiments on environmental samples exposed to different treatments and measured the CO_2_ production from oil degradation via reverse stable isotope labeling (RSIL) method for the calculation of oil degradation rates and analyzed the change in the microbial community composition via 16S rRNA gene amplicon sequencing. Additionally, visual and microscopic observations, pH measurements were collected from the microcosm incubations over the 81-day period. The findings from this work contribute to the understanding of aerobic microbial oil degradation in adapted and naïve freshwaters. Moreover, it expands the knowledge of the application of hydrocarbon adapted seedbanks to anthropogenic oil slicks in order to speed up degradation, including the identification of key players for the degradation of naturally abundant hydrocarbons in freshwaters. Furthermore, this knowledge could be used for the remediation of oil-polluted freshwater environments.

## Material and Methods

### Oil and water sampling

Heavy oil was sampled from the Hänigsen Teerkuhle (tar pit, 52°29’38.6“N 10°05’28.3”E). At this site, around 10 L oil per year continuously rise from a ∼1 km-deep subsurface reservoir into a pit built around the natural seep, first documented in 1546 (27). An oil slick on the groundwater, which fills the pit, was sampled by sideways immersion of a sterile 2 L Schott flask. Water beneath the oil was sampled by using a strong fan to push the oil aside, allowing collection with a sterile 2 L Schott flask without excessive oil slick contamination. Since the tar pit’s subsurface oil reservoir remained largely untouched by human activity, the microbial community in the oil is expected to be naturally adapted to this habitat. The same likely applies to the water beneath the heavy oil. The oil from the tar pit is highly viscous (kinematic viscosity of 895 mm²/s at 40°C) and has a density of 0.961 kg/L at 15°C. Both parameters indicate a high degree of oil degradation which is typical for heavy oil. The heavy oil was stored at 4 °C until used.

Light oil was provided from the oil and gas company Aker BP ASA and was obtained in 2016 from a Norwegian offshore oil reservoir from 3700 m depth (28). The light oil has a density of 847.6 kg/m³ at 15°C and API 37 (28). It is much less viscous compared to the heavy oil (API = 15.7). The light oil was stored in the dark at room temperature in the dark until used. Both oils were not autoclaved to preserve its chemical composition and to include the existing microorganisms that potentially contribute to oil degradation.

Naïve water was taken from the Aller river (29323 Wietze, 52°39’56“N 9°48’34”E), ca. 1 m away from the riverbank by immersion of a sterile 2 L Schott flasks and the following parameters were measured *in situ*: pH: 7; conductivity: 0.66 mS/cm; temperature: 17 °C. This river water is considered naïve to oil, because it is not chronically exposed to oil, thereby potentially hosting microorganisms that are not adapted to the utilization of carbon sources that are characteristic for oil. The water was stored at 4 °C until used.

### Microcosm setup and sampling

Microcosms were set up in sterile 250 mL Schott flasks. The adapted (from the tar pit) and naïve (river Aller) waters were separately filtered through a 10 µm filter (Acrodise PSF syringe Filter, Pall) to exclude protists. From this filtered water, 100 mL were added to sterile 250 mL Schott flasks with sterile serological pipettes. To each microcosm 1 mL of oil (either heavy oil or light oil or a 1:1 mix) was added by slow pipetting with a piston pipette, to ensure that the viscous oil will have an accurate volume. Next, the flasks were sealed with butyl stoppers (Ochs Laborbedarf, Germany). The butyl stoppers were heat treated beforehand to remove excess plasticizers (2x 30 min in boiling water) and autoclaved before usage. Headspace of the microcosms remained ambient air, the headspace volume of all microcosms was 210 mL and increased with every sampling. As control, one treatment (A2) was incubated in an anoxic atmosphere, where the headspace was flushed with pure N_2_ for 5 min before and after sealing the bottle with butyl stoppers. After the microcosms were sealed, sterile NaHCO_3_-Buffer was added anoxically to provide a known ^13^C background for RSIL. The applied hydrogen carbonate buffer was ^13^C-labeled with x(^13^C) = 10 atom % as a mixture of regular NaHCO_3_ x(^12^C) = 1.11% (Carl Roth, Germany) and ^13^C-labeled NaHCO_3_ x(^13^C) = 98% (Sigma-Aldrich, MO, U.S.A.) with final concentration of 10 mM. No additional carbon sources, nutrients or supplements were added, so that water and oil were the only carbon sources. Afterwards, ^13^C-label was added anoxically. A total of 28 microcosms were produced to yield seven distinct conditions for oil degrading microbial consortia, which included each oil type (light, heavy and 1:1-oil mix of both) combined with each water type (naïve and adapted) and an anoxic setup with adapted water and heavy oil. The microcosms were incubated in the dark at 20 °C without shaking. Microcosm treatments were categorized into two groups: “adapted habitat”, which included water from the oil-adapted tar pit, and “naïve habitat”, which included oil-unexposed water from the river Aller. Each test condition was set up in biological triplicates with one additional autoclaved control (121°C, 20 min), serving as negative controls. Following conditions were tested: Adapted water + heavy oil (A1), anoxic: adapted water + heavy oil (A2), adapted water + light oil (A3), adapted water + 1:1 mixed oils (M7), naïve water + heavy oil (N4), naïve water + light oil (N5), naïve water + 1:1 mixed oils (N6, supplementary data S1). All microcosms were sampled under sterile and anoxic conditions to avoid the introduction of other organisms or CO_2_. Syringes (3 mL, 5.0 grade; sterile Hungate needle) were flushed three times with sterile N_2_-gas before piercing the butyl stoppers. For each sampled microcosm, 5mL aqueous phase were taken, from which 3 mL were filtered (0.2 µm, Filtropur S0.2, Sarstedt AG&CoKG) before 2x 500 µL were put into prepared 12 mL Labco exertainer vials (Labco Limited, UK) for isotopic CO_2_ measurement via reverse stable isotope labeling (RSIL) method. Labco exertainer vials were preamended with 50 µL of 85% phosphoric acid, closed with screw caps containing butyl rubber septa, and flushed with CO_2_-free synthetic air (6.0 grade; Air Liquide, Germany) before usage. The remaining 2 mL were put unfiltered into a DNAse free reaction tube (Sarstedt AG & Co. KG, Nümbrecht, Germany) for 16S rRNA gene amplicon sequencing and from there distributed to new reaction tubes for cell counting (see supplement) and immediate pH measurement (Mettler Toledo FiveEasy F20, 0.3 mL). Samples taken for 16S rRNA gene amplicon sequencing were kept on ice during the sampling and were frozen at −70 °C until use. Initially, the sampling was performed after 0, 2, 5, 10 days of incubation and afterwards, every 7 (± 1) days until 81 days of incubation. Cell morphologies were occasionally observed with a Nikon Eclipse Ni (Nikon Europe B.V.) with 20x and 40x oculars. The software NIS-Elements was utilized for image processing.

### Assessing oil mineralization rates

Microbial oil degradation is associated with a decrease in pH (29). For this purpose, pH values were determined with a pH electrode (Mettler Toledo FiveEasy F20, InLab Flex-Micro) during microcosm sampling. The error of the employed electrode was ± 0.3. The electrode was calibrated with three standard solutions with the following pH: 4.006; 6.865 and 9.180 (Xylem Analytics Germany GmbH WTW) prior to the measurement. The R^2^ was at least 0.9, otherwise the electrode was recalibrated.

To determine the CO_2_ production from microbial oil degradation, reverse stable isotope labeling (RSIL) method was applied. Water phase of all 28 microcosms was sampled 12 times over 81 days to assess the microbial degradation rates of microbial communities fed with either heavy oil, light oil, or both oils mixed (1:1). All samples for the reverse stable isotope labeling measurement were taken in technical duplicates under anoxic conditions with N_2_-gas flushed syringes and filtered (0.2 µm, Sarstedt AG & Co. KG, Nümbrecht, Germany) before 500 µL were put into prepared 12 mL Labco exertainer vials, as described above. Samples were analyzed with a Delta Ray CO_2_ Isotope Ratio Infrared Spectrometer (Thermo Fisher Scientific, MA, U.S.A) with Universal Reference Interface Connect for measuring carbon isotope compositions of produced CO_2_. CO_2_-free synthetic air (Air Liquide, Germany) was used as carrier gas. CO_2_ in synthetic air at 414.2 ppm (Air Liquide, Germany) was used for CO_2_ concentration calibration. CO_2_ reference gases used for calibration of carbon isotope ratios had δ^13^C values of −9.7‰ (Thermo Fisher, Bremen, Germany) and x(^13^C) = 10% (Sigma-Aldrich, Taufkirchen, Germany). Pure CO_2_ gas with x(^13^C) = 10% was used as working reference gas. The CO_2_ concentration for reference and sample gas entering the analyzer was set to 380 ppm for optimal precision. Each sample was measured for 5 min and the obtained δ^13^C values were averaged. The stable carbon isotope data were received as delta values and converted into isotope-amount fraction and total produced CO_2_ in mM as published before (30–33). The mean of the technical duplicates per microcosm was used for all calculations to determine total produced CO_2_ per biological replicate. CO₂ concentrations measured in sterilized controls were subtracted from experimental values to correct for potential measurement artifacts. Next, the mean of the biological triplicates was determined and plotted with the standard deviation as error bars against the incubation time in days. For the calculation of the CO_2_ production rates per day, the linear part of the plot was used (between days 0 and 52). The slope of this plot equates to the mean CO_2_ production rate expressed in mM day^-1^. To compare oil degradation rates between two treatment groups, a one-tailed independent t-test assuming equal variances was performed. Replicate values from each condition were used as input. A *p*-value of < 0.05 was considered statistically significant. Details of the mass balance, used to assess oxygen consumption and to quantify the extent of oil degradation, are provided in the supplementary text.

### Microbial community composition analysis via 16S rRNA gene amplicon sequencing

DNA extractions from each microcosm was performed after 153 days of incubation, using the DNeasy PowerLyser Power Soil Kit (Qiagen GmbH, Germany) following the manufacturer’s instructions with the following modifications: Steps 14 and 15 were repeated three times and each time solution C5 was added in quantities of 250 µL. In step 15, centrifugation time was increased to 2 min and air dried after the third centrifugation step. Steps 17 and 18 were repeated two times. Each time 25 µL solution C6 was added and incubated at room temperature for 2 min. The centrifugation time for step 18 was increased to two min. Extracted DNA and PCR products were purified (MagSi NGSprep Plus magnetic beads, Beckman Coulter GmbH, Germany) and processed according to the Illumina protocol “16S Metagenomic Sequencing Library Preparation” (Document No. 15044223). Amplicon-based PCR was performed to amplify the V4 region of the 16S rRNA gene using universal standard primer set S-D-Arch-0519-a-S-15 and S-D-Bact-0785-a-A-21 (34). Each 25 µL PCR reaction contained 12.5 µl polymerase 2x Kapa Ready to use mix (Roche), 1 µL of each primer 0.2 mM, 3 µL template DNA and 7.5 µL PCR-water. Thermal cycling conditions were as follows: initial denaturation at 95 °C for 3 min, followed by 38 cycles of 95 °C for 30 s, 55 °C for 45 s, and 72 °C for 1 min, with a final extension at 72 °C for 5 min. Illumina indices and adapters were added in a final PCR step using the Nextera XT DNA Library Preparation Kit v2, Set D (FC-131-2004; Illumina, Munich, Germany). All PCR products were checked with agarose-gel-electrophoresis using a 1.5 % agarose gel in 1x TRIS-Acetat-EDTA buffer (Merck KGaA, Darmstadt). The gel was stained with 1 µL SYBRR Save dye and 5 μL PCR-product was mixed with 2 µL loading dye, whereof 5 µL were loaded onto the gel. Electrophoresis was performed at 100 V for 45 min (Amersham Biosciences, Hoefer HE33) in the dark. For visualization of the separated DNA bands, a gel documentation system (BioRad) was used in UV mode with the software Quantity one 4.6.3. Promega G5711 1 kb DNA ladder was used. The DNA concentration of each sample was normalized to 10 ng µL^-1^ and pooled into one ready-to-load library. During the whole library preparation negative DNA-extraction and PCR-controls as well as a commercial whole cell positive control was applied (ZymoBIOMICS® Microbial Community D6300, USA). Due to samples resources and technical issues during DNA-extraction, high quality DNA was only received for beginning of the experiment from incubations adapted water + heavy oil (A1), adapted water + heavy oil (anoxic, A2) and adapted water + light oil (A3). No DNA was extractable from naïve water + heavy oil (N4), naïve water + light oil (N5), naïve water + 1:1 mixed oils (M6) or from adapted water + 1:1 mixed oils (M7). Here DNA was only received after 153 days of incubation. Illumina MiSeq sequencing with V2 500 cycle flow cell was performed by the external company CeGaT GmbH (Tübingen, Germany). Demultiplexing and adapter trimming was performed by CeGaT GmbH. Resulting sequences were analyzed R-packages DADA2 pipeline v 1.30. 0 (35) and the R package phyloseq v1.52.0 (36). Forward and reverse reads were trimmed to 240 and 160 base pairs with following conditions: filterAndTrim(fnFs, filtFs, fnRs, filtRs, truncLen=c(240,160), trimLeft=c(14,21), maxN=0, maxEE=c(8,8), truncQ=2, rm.phix=TRUE, compress=TRUE, multithread=FALSE) prior to merging and chimera removal. Taxonomic assignment was performed using the SILVA database SSU138.2 database (37), and sequences were clustered into Amplicon Sequence Variants (ASVs) using DADA2. To minimize the influence of rare organisms and potential sequencing artifacts, ASVs were removed if they had fewer than 10 reads or where taxonomy was classified as “NA”, Kingdom=”NA”, chloroplasts and mitochondria.

Additional to simple literature research, the full standard PICRUSt 2.0 (Phylogenetic Investigation of Communities by Reconstruction of Unobserved States (38) version 2.6.2 was applied under default settings for the comparison of potential metabolic differences between the samples, with special focus on hydrocarbon degradation.

Raw sequencing data are available under the NCBI database BioProject PRJNA1365740.

### Determination of the initial dissolved organic carbon concentration in the environmental samples

Heterotrophic microorganisms can feed of a wide variety of carbon sources. Though, this study focused on oil degradation which necessitates that no additional carbon sources are available for heterotrophic growth. The dissolved organic carbon content of the adapted water and naïve water was determined to account for potential growth from dissolved organic carbon in the CO_2_ production experiments. Dissolved organic carbon was analyzed with a TOC-L total organic carbon analyzer (Shimadzu, Duisburg, Germany). For sample preparation, triplicate samples of 20 mL were filtered through 0.45 μm cellulose-acetate filters (Sarstedt AG & Co. KG, Nümbrecht, Germany). For the determination of the non-purgeable organic carbon, 10 mL of the sample were transferred into open glass vials for the TOC-L autosampler (Shimadzu, Duisburg, Germany). The whole system was washed twice with the sample to avoid carry-over between the individual samples, before the measurement. The non-purgeable dissolved organic carbon was determined by purging the sample for 3 min with synthetic air (80 mL·min^−1^) inside the syringe of the autosampler unit. Afterwards, 50 μL of the sample were injected into an oxidation tube filled with platinum coated ceramic particles at 720 °C. The CO_2_ formed during the oxidation was measured by infrared spectroscopy. A calibration was performed with a certified TOC reference standard (Sigma Aldrich, 1000 ± 10 mg·L^−1^ TOC) that was prepared to 0, 2, 5, 10, 15, 20, 25 and 50 mg·L^−1^.

## Results

We have performed a literature survey clearly highlighting the imbalance in research focus between marine and freshwater oil slicks. Marine systems dominate the published work since the year 1900, with 7,822 studies compared to 1,549 on freshwater oil slicks, indicating that freshwater oil contamination is comparatively understudied. When considering microbial investigations, this disparity is even more pronounced, only 275 studies focus on microbial processes in freshwater slicks versus 1,491 in marine environments. These numbers underscore the need for targeted research on microbial dynamics in freshwater oil-contaminated systems, which remain a relatively small and underexplored subset of the literature. To address this gap, we investigated the potential of using naturally adapted microorganisms to oil as a carbon source from a crude oil seep, the tar pit in Hänigsen, with the goal to accelerate the degradation of anthropogenic oil slicks in previously oil-naïve aquatic environments.

### Oil snow formation and pH decrease in the microcosms indicated oil degradation

The first hints for oil degradation were observed after about 30 incubation days by continuous acidification and visual changes to the microcosms. The formation of oil snow became clearly visible after about 70 days of incubation, particularly evident in the heavy oil treatments (Figure 1). In the light oil incubations, the oil degraded into much finer particles and these particles did not precipitate but rather accumulated in the oil slick on the water surface and became visible after shaking. Such oil snow formation was not observed in any of the autoclaved controls, where the medium stayed clear, the oil remained as an intact film on the water surface without oil snow formation. When oil snow was investigated under the microscope, it showed a heterogenous structure with differently shaped, sized and colored oil droplets with either clear rims in the light oil or very irregular rims in the light oil, indicating degradation processes.

**Figure 1.**
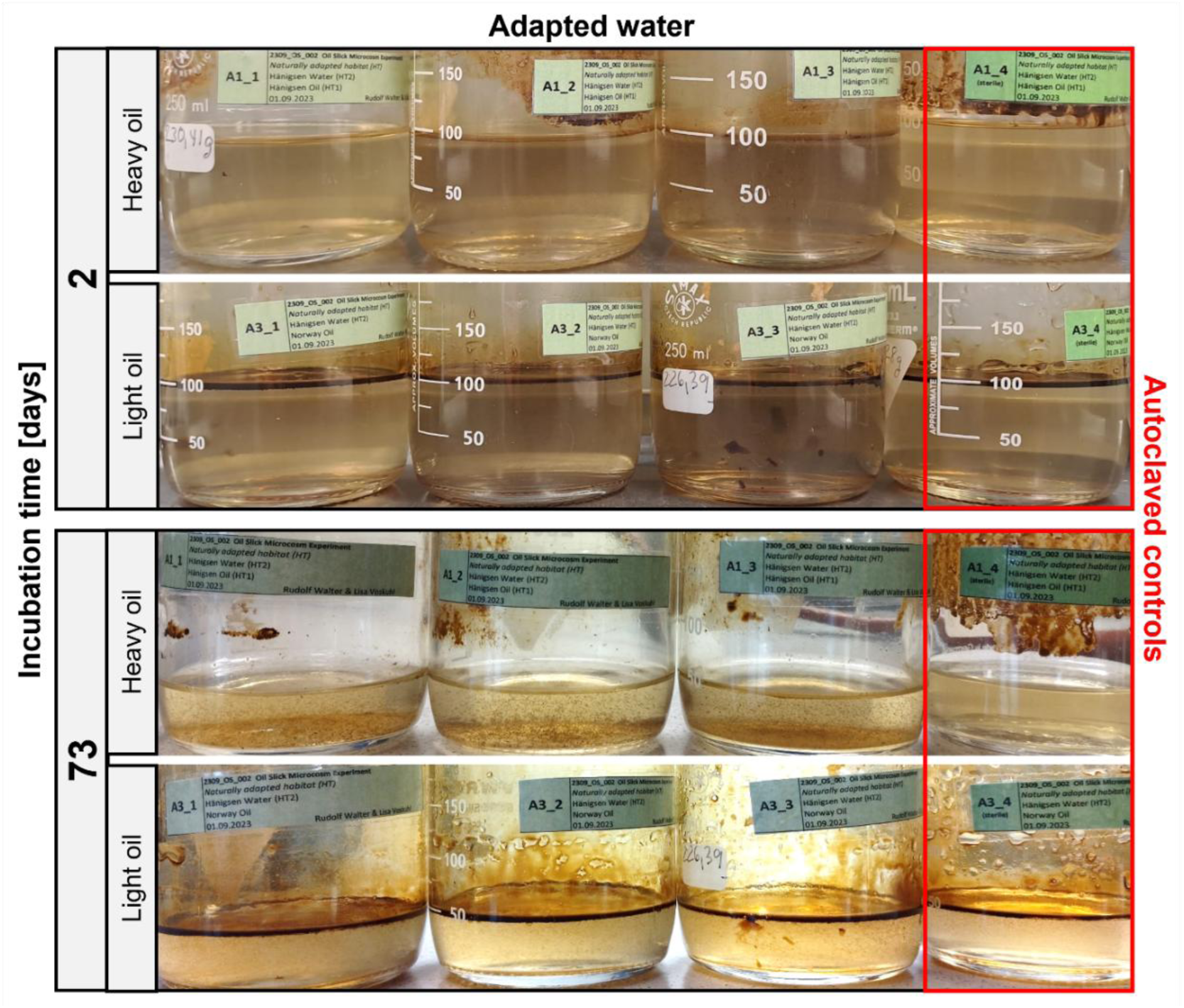
Microcosm incubations with adapted water with heavy and light oil addition 2 days and 73 days after inoculation. After 73 days oil snow was clearly visible.

The initial average pH in each microcosm was 8.5 ± 0.2 standard deviation. As expected, the smallest pH change was observed for the anoxic incubations (ΔpH=0.36 ± 0.16, n=3) and the sterilized controls (ΔpH=0.38 ± 0.13, n=5), but this change was so small that it fell within the error range of the utilized pH electrode (± 0.3) and is therefore regarded as not meaningful. A strong pH decrease was observed for all other setups, ranging from ΔpH=0.74 ± 0.16 in naïve light oil incubations and ΔpH=1.12 ± 0.21 in the adapted heavy oil incubations over ΔpH=1.17 ± 0.07 in the naïve heavy oil incubations and ΔpH=1.19 ± 0.16 in in mixed oil incubations with adapted water and light oil addition. Largest pH changes were observed in the adapted water microcosms treated with light oil ΔpH=1.26 ± 0.04 and the naïve water with light oil ΔpH=1.42 ± 0.11 (Figure 2, Data S2).

**Figure 2.**
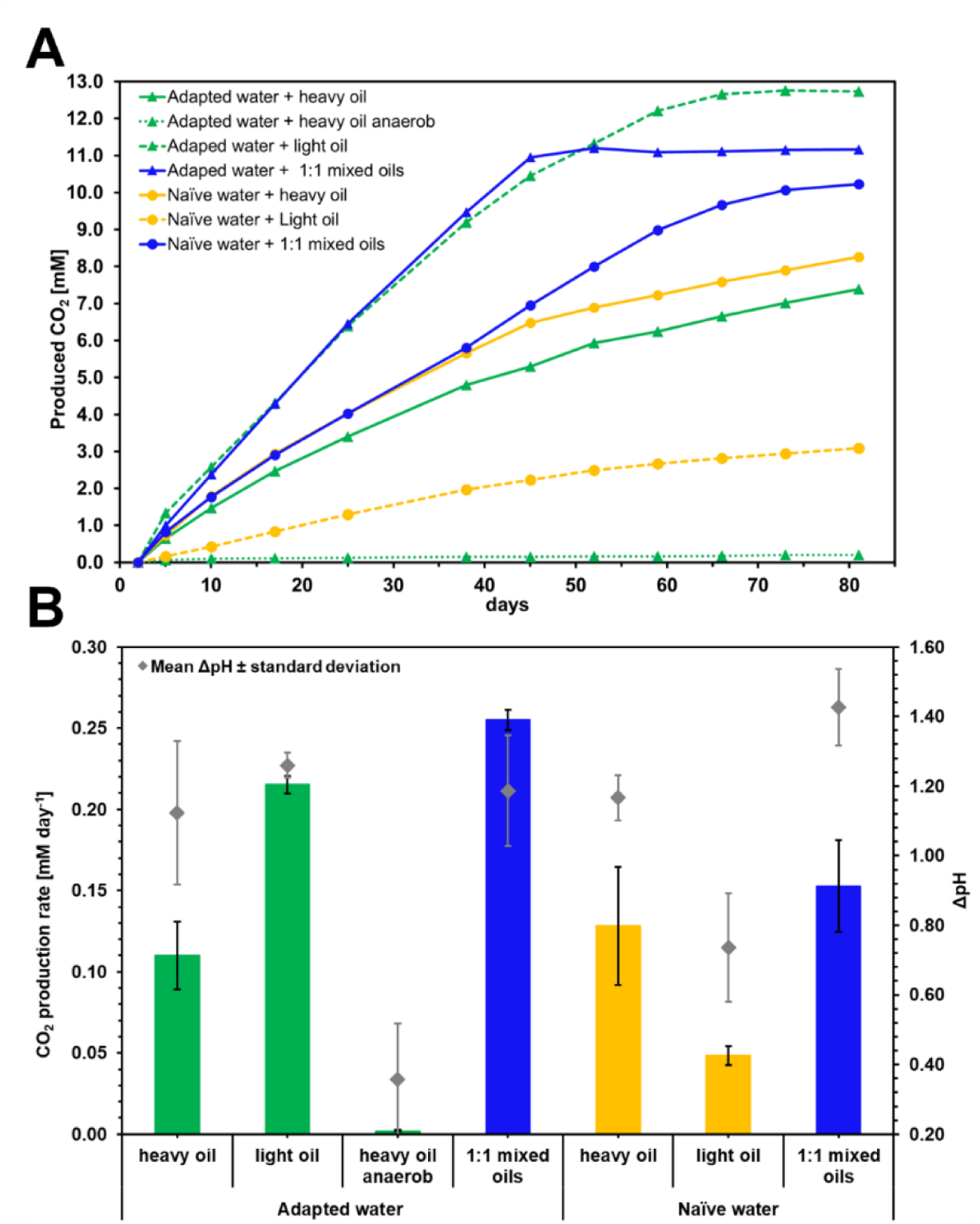
Produced CO2 from oil degradation **A)** Mean produced CO2 over time from three biological replicates in the different treatments of adapted waters from the tar pit and naïve Aller water incubated with heavy oil, light oil and a 1:1 mix of both oils with oil as main carbon source. Error bars are not indicated for better readability, for standard deviations see (Data S4). To account for measurement artifacts, CO2 concentrations from sterilized controls were subtracted from the experimental data. **B)** Bar graph comparison of mean CO2 production rates, categorized in naturally adapted habitat (green), naïve habitat (yellow), mixed oils with adapted *and* naïve water (blue). The linear part from CO2 production curves (Figure 2A) were used until day 52 to obtain the CO2 production rate in mM/day. Error bars indicate the standard deviation between the three biological replicates. Rhombuses display the mean change of the pH (ΔpH, n=3) and standard deviations between biological replicates. The trend of pH change during incubation shows the same pattern as CO2 production rates.

### Highest oil biodegradation rates by microorganisms adapted to natural oil seeps

The dissolved organic carbon content in naïve water from the Aller river was determined 14.42 mg L^-1^. The CO_2_ production resulting from microbial oil degradation was monitored over 81 days with the reverse stable isotope labeling method. Microbial oil degradation rates were calculated from the slope of the first 52 days of the CO_2_ production curve of the different microcosm treatment, where the CO_2_ increase was linear and not yet reaching a plateau (Figure 2A). As expected, the anoxic treatments (Figure 2A, dotted green line, triangles), serving as an additional control, showed with 0.002 ± 0.0005 mM produced CO_2_ per day the significantly lowest CO_2_-production rate (all *p* < 0.002, t-test) as no electron acceptor, such as oxygen, was available. The microcosms with adapted water and heavy oil (Figure 2A, solid green line, triangles) and naïve water with heavy oil (Figure 2A, solid yellow line, circles) showed comparable CO_2_ production curves and comparable CO_2_ production rates (0.11 and 0.13 mM day^-1^, respectively) showing no significant differences between them (*p* > 0.248, t-test). This indicated that the oil degrading microorganism were sitting in water inclusions in the heavy oil, rather than originating from the aqueous phase of the naïve water. The CO_2_ production curve (*p* = 0.014) and rate (0.22 ± 0.005 mM day^-1^, *p* = 0.001) from adapted water with light oil (Figure 2A, dashed green line, triangles) were significantly higher compared to the adapted water treated with heavy oil. This demonstrated that the adapted water contains an oil degrading microbiome that can gain more energy and faster from light oil rather than from the heavy oil, revealing that the microbiome is adapted but not restricted to the consumption of the heavy oil, it is used to live in. Naïve water incubated with light oil Figure 2A, dashed yellow line, circles) showed the lowest CO_2_ production curve and with 0.048 ± 0.006 mM day^-1^ the lowest CO_2_ production rate along all oxic treatments, being significantly different to all other treatments (*p* < 0.01, Figure 2B), implying that the use of a microbial seedbank could successfully increase oil degradation rates and that there is a potential for additional degradation through the seed bank, because the naïve light oil shows slower degradation rates. Indeed, the oil degradation could be accelerated significantly by amending light oil with the microbial seedbank from the heavy oil, containing a well-adapted oil degrading microbial community. The microcosms with adapted water and the 1:1 mixed oils (Figure 2A, solid blue line, triangles) produced comparable CO_2_ concentrations to the adapted water with light oil, but the adapted water with 1:1 mixed oils was slightly faster but significantly higher (*p* = 0.001) with a CO_2_ production rate of 0.26 ± 0.006 mM day^-1^ which plateaued already after 52 days, while the CO_2_ production increased further with light oil (0.22 ± 0.005 mM day^-1^), as here more degradable oil was provided. The naïve water with 1:1 mixed oils (Figure 2A and B, solid blue line, circles) showed a significantly higher CO_2_ production rate of 0.15 ± 0.028 mM CO^2^ day^-1^ compared to the naïve light oil (*p* = 0.002, Figure 2B) but not to the naïve heavy oil incubation (*p* = 0.203), indicating once again, that most likely the microbes from the heavy oil were responsible for the faster oil degradation.

After 52 days of incubation, the naïve water with 1:1 mixed oils reached much higher degradation rates in comparison with the naïve water with heavy oil, but lower than the adapted water with 1:1 mixed oils. Overall, the data demonstrated that in adapted water with light oil, the biodegradation is two times faster than with heavy oil. Microorganisms originating from the natural oil seep were able to degrade oil when introduced into freshly contaminated, oil-naïve water. While here biodegradation occurred for both oil types, the heavy oil in the naïve water was degraded approximately 2.6 times faster than the light oil, again showing that the adapted microbial community sits closely attached in the heavy oil film, because here the rate was similar to the adapted water treated with heavy oil. In the naturally adapted water, degradation of 1:1 mixed oils was approximately 1.7 times faster compared to the naïve water with mixed oils. Adding heavy oil as seedbank into contaminated naïve water (1:1 oil mix) enhanced degradation rates by about 3.2 times compared to the naïve water only with light oil. The CO_2_ production of the adapted water with light oil (Figure 2A, dashed green graph, triangles), the adapted water with mixed oils microcosms (Figure 2A, blue graph, triangles) and naïve water with mixed oils (Figure 2A, blue graph, circle), plateaued at 12.8 mM, 11.2 mM, and 10 mM produced CO_2_. At 12.8 mM we could calculate an oxygen limitation in the closed microcosm flasks. For the degradation of 1 g oil 2.4 L oxygen are required, however only 42 mL O_2_ were initially available in the headspace, resulting in a theoretically maximal degradation of 17.5 mg. This amount was almost reached in the microcosms, because at the plateau in graph Figure 2A ∼16.8 mg of oil was degraded with the consumption of ∼1.8 mmol O_2_ (see calculation in supplement text). In case of ongoing anaerobic biodegradation, CO_2_ development would be barely noticeable since the anoxic incubation (Figure 2A, green graph, circle) produced only 0.2 mM CO_2_ in total, which is only 1.7% of the produced CO_2_ in aerobic incubations.

### Microbial communities differed based on the treatment

16S rRNA gene amplicon sequencing of environmental samples from the naïve Aller water, the adapted Hänigsen tar pit water, and all microcosm incubations after 153 days demonstrated that the microbial communities differed between environmental and microcosm samples from beginning on (Figure 3, Figure S6), displaying a strong inoculation effect. Compositional ordination of microbial community compositions based on Bray–Curtis dissimilarity showed that the microbial communities in the autoclaved microcosms (sterile controls, squares) contained mostly different community compositions compared to the incubated microcosms (triangles), with exception for adapted water treated with light oil (Figure 3, A3, red), where these differences remained low, demonstrating that microbial activity (CO_2_ production) occurred together with growth in the microcosms. Most different from all other samples were the anoxic incubated microcosms from adapted water with heavy oil addition (A2, red), demonstrating that the Hänigsen tar pit hosts aerobic and anaerobic microorganisms capable for oil degradation. As these communities were completely different to the oxic incubations, this showed that after reaching oxygen limitation in the microcosms (containing the adapted water and light oil (A3), adapted water with 1:1 mixed oils (M7), and naïve water with 1:1 mixed oils (M6)), the microbial communities did not change drastically towards an anaerobic community. Microbial communities from adapted water incubated with heavy oil (A1) were most different to the other microbial communities with exception for one naïve water plus heavy oil microcosm (N4) and the sterile (N4), suggesting that the microbiome in the tar pit is unique and cannot be completely displayed by only the water or only the oil.

**Figure 3.**
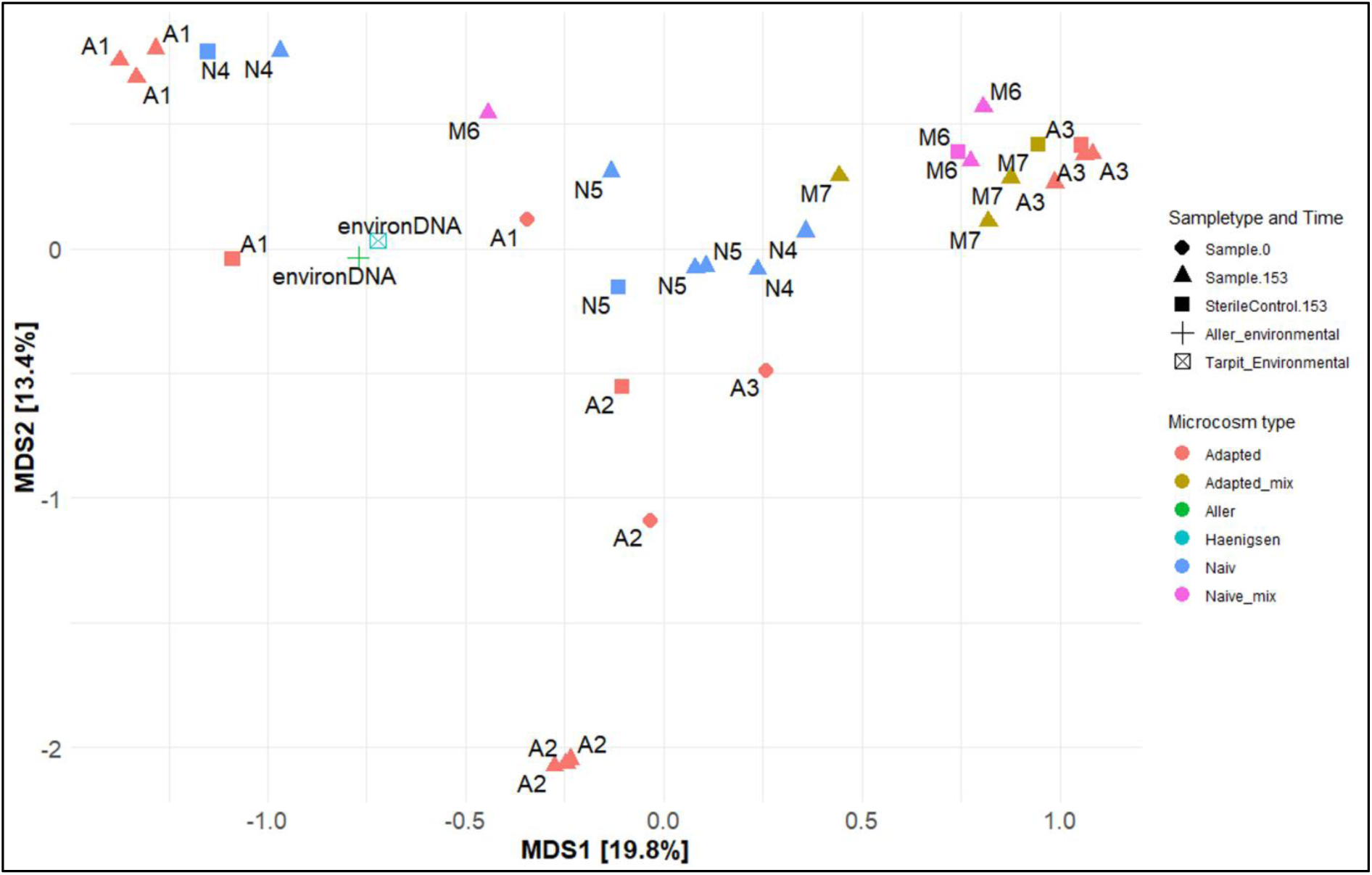
Beta diversity: ordination of microbial community compositions based on Bray–Curtis dissimilarity. Ordination was performed on relative abundance data transformed at the unique taxonomic rank. The plot displays the first two ordination axes (MDS and MDS2), showing differences in community structure among samples. Points are colored and filled according to Microcosm type, while point shapes represent sample type and time of DNA extraction. Sample labels indicate individual experimental units: A1: adapted water + heavy oil, A2: adapted water + heavy oil, anoxic, A3: adapted water + light oil, M7: adapted water + 1:1 mixed oils, N4: naïve water + heavy oil, N5: naïve water + light oil, M6: naïve water + 1:1 mixed oils (Table S1). The analysis was conducted using Bray–Curtis distances and an automatic ordination method.

Otherwise, the other microcosms with adapted tar pit water (amended with light oil (A3) or with the 1:1 mixed oil (M7)) would be more similar to the adapted water incubated with heavy oil (A1). Microcosm microbial communities from naïve water treated with heavy oil (N4) and with light oil (N5) were more similar to each other building a cluster, so the oil type was not affecting the total microbial community strongly (Figure S5). And microcosms with mixed oils in adapted tar pit water (M7) and naïve river Aller water (M6) built a cluster with adapted water treated with light oil (A3). This suggested that some microbes are sitting in the heavy oil, most likely in water inclusion and contribute to the fast oil degrading microbial communities. Despite some incubations already reaching oxygen limitation after around 70-80 days of incubation, the microbial communities after 153 days were different to the anoxic incubated microcosms. Indicating more that the microbial communities did not change too much since beginning of the limitation, as observed also for the cultures that were immediately incubated anoxic (A2). This suggested that anaerobic microbial communities that established after electron acceptor limitation are different from the immediate anaerobic microcosm incubation (A2).

All investigated treatments shared a large core of 1215 ASVs with abundances above 0.01 % (sterile controls excluded, data S10). Most unique ASVs (386) were found in the light oil treated samples. In the heavy oil treated samples 291 unique ASV were identified and 190 ASVs were found uniquely in the mixed oils. Some of the unique ASV in the mixed oils treatments were also present in other samples but showed an abundance below 0.01 %.

Most ASVs (352) were shared between the heavy oil and light oil treatments indicating that also lots of oil degrading microorganism were originating from the waters.

### Metabolic predictions did not fit to the measured CO_2_ production rates

In order to identify potential oil degrading microorganisms, we applied the tool PICRUSt2, genes for central metabolisms (data S6) were equally distributed along all samples while genes for chosen pathways that are related to hydrocarbon degradation showed sample specific differences (Figure 4, Figure S6), that however showed a different picture than our measured CO_2_-production rates (Figure 2B). Based on 16S rRNA gene sequence abundances, highest oil degradation potential was estimated for the naïve water sample with light oil. However, CO_2_ measurements revealed that these were the microcosms with the lowest oil degradation. Genes for aromatic compound degradation contributed max 1.07% in the naïve water treated with light oil over all genes. The degradation potential of aromatic compounds was significantly lowest in anaerobic cultures (t-test: <0.0001). Toluene degradation potential was always higher in heavy oil treated set ups (Figure 4, data S7). By looking only at two marker genes for aerobic activation steps in hydrocarbon degradation, alkane 1-monooxygenase (alkB1_2, alkM) and long-chain alkane monooxygenase (ladA, data S8) also did not align with the measured CO_2_ production rates (Figure 2B).

**Figure 4.**
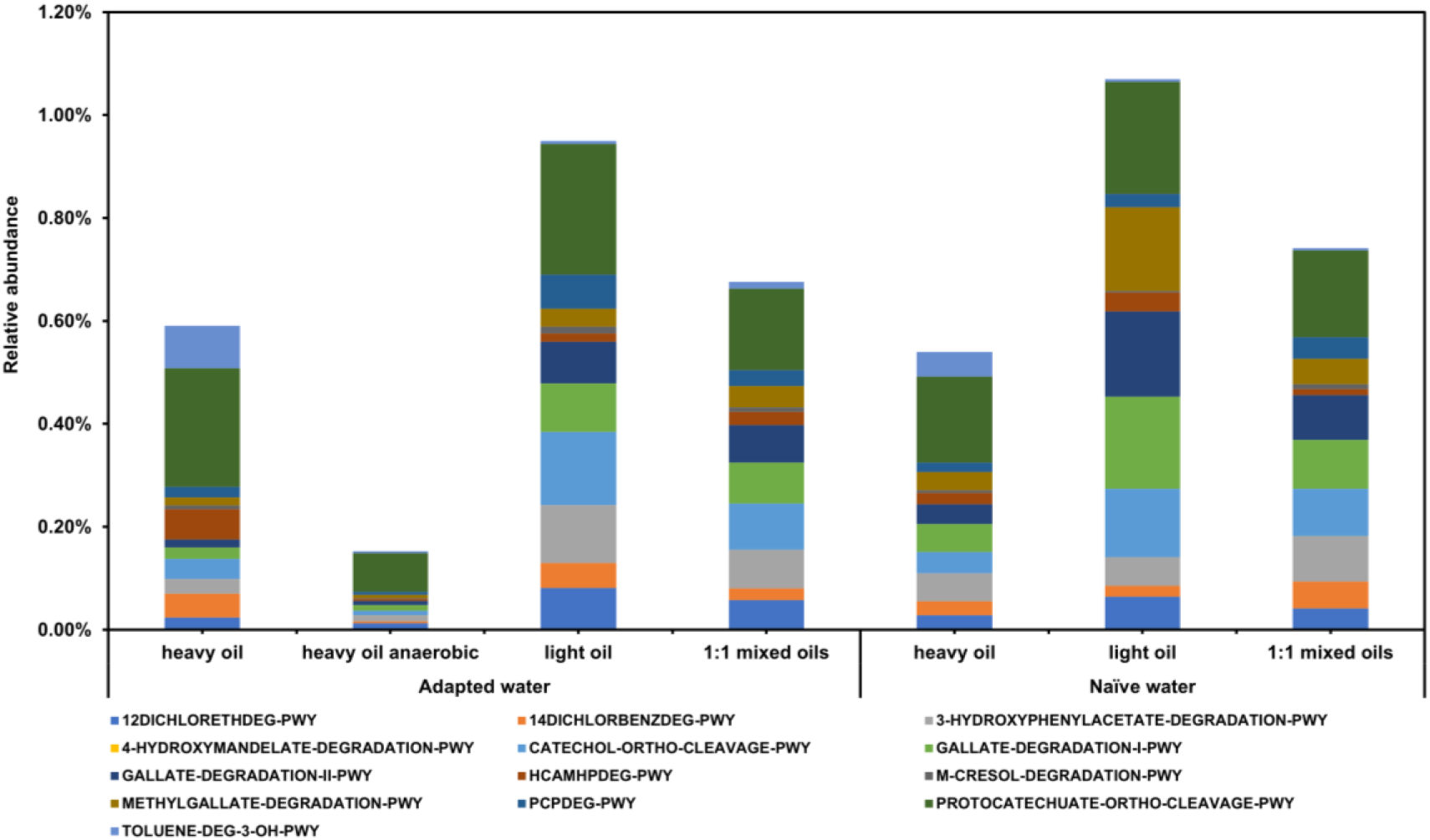
Mean abundance of potential hydrocarbon degradation pathways in the microbial communities after 153 days of incubation estimated with PICRUSt2.

This demonstrated that such predictive tools based on 16S rRNA gene amplicon data should never be applied without laboratory experiments, as this could otherwise lead to wrong conclusions. In total PICRUSt2 helped to identify 1080 potential oil degrading ASVs in our samples of which 339 where unique taxa potentially oil degrading. Most identified taxa are known from literature to be capable of oil degradation but were quite low abundant in the samples: *Nevskia* (ASVs: 1, 3175, 3946)*, Acinetobacter* (ASVs: 1098, 1445, 1992, 2422, 2547, 3268, 3545, 4918)*, Burkholderia-Caballeronia-Paraburkholderia* (ASVs: 16, 395, 531, 534, 696, 728, 1018, 1146, 1660, 1953, 2021, 3984)*, Amaricoccus* (ASVs: 427, 527, 2759, 2844, 3162, 4014)*, Aquabacterium* (ASVs: 579, 1133, 1939, 2830)*, Aquisphaera* (ASVs: 1765, 2511, 3217, 3508), Candidatus *Koribacter* (ASVs: 2421, 3826, 4117, 4165, 4282, 4839), Candidatus *Solibacter* (ASVs: 4364, 5050, 5124, 5732)*, Solirubrobacterales* (ASVs: 178, 503, 747, 833, plus 57 more)*, Blastocatellia* (28 ASVs, data S9, eleven genes were chosen for hydrocarbon degradation)*. Nevskia* is capable of oil degradation (40), was mainly found in the adapted water samples at the beginning of our experiment (T0) and vanished completely over incubation time, so that this organism was unlikely one of the active oil degraders. A review article summarized that many *Acinetobacter* species are commonly isolated from waste water treatment plants and sewage and that they are reported to be capable of crude oil degradation by making use of aliphatic and aromatic components. *Aquabacterium olei* was isolated from oil-contaminated soil in Korea and was described as an oil-degrading bacterium (41). The most abundant ASV along all samples was ASV26 *Parablastomonas* originating from the naïve water, its direct role in hydrocarbon degradation remains unconfirmed, but members of its family *Sphingomonadaceae* have been detected in contaminated soils (42). ASV7 *KCM-B-112* (family *Acidithiobacillaceae*) was second most abundant ASV along all samples and most abundant in the adapted water with heavy oil, this organism has been repeatedly detected in hydrocarbon-impacted soils and oil sands tailings (43, 44) and is therefore a likely candidate as a biomarker of oil contaminated sites.

## Discussion

This study demonstrated that microorganism living in an oil slick of a natural oil seep are highly adapted to hydrocarbon degradation of heavy and light oil, whereof light oil is degraded two times faster. This oil slick itself, together with the attached microbes and also the adapted groundwater, can be applied as a seedbank for enhanced hydrocarbon biodegradation in another freshwater body that has never been in contact to oil before. The presented results provide a quantified perspective on the application of microbial seedbanks and their use for accelerating oil degradation in case of anthropogenic oil spills.

### Oil snow formation and pH changes reveal ongoing degradation processes

The observation of the development of oil flakes during oil degradation is a well-known phenomenon from marine environments and is often referred to as (marine) oil snow (45, 46). Upon oil exposure, bacteria release extracellular polymeric substances to which cells and other particles attach, most likely as a defense mechanism (protection from toxic hydrocarbons) or as an attachment mechanism to facilitate oil decomposition for energy production (47, 48). These extracellular polymeric substances are precursors of oil snow, which is known to accumulate high molecular weight compounds like polyaromatic hydrocarbons (49). Oil snow exerts a wide range of impact on flora either directly or indirectly. It can create suboxic and anoxic conditions (50), immunotoxicity through gills and can lead to physical smothering of animal tissues or egg surfaces (51). Furthermore, bioaccumulation of petroleum compounds in the planktonic food web ultimately results in exposure and biomagnification of higher trophic-level organisms (52). The acidification due to oil biodegradation is well known and is related to the production and accumulation of naphthenic acids (29). Our measured trends in the decrease of ΔpH values fitted the measured CO_2_ production rates well (Figure 2B), revealing that the change in pH gives already fast and low-cost impressions on potential degradation processes. Measuring the pH value during oil degradation experiments could avoid expensive and complex analytical sample preparations and help to find the right time point to perform analytical experiments.

### Discrepancy between metabolic predictions and CO_2_ measurements

As we could visually observe hydrocarbon oil degradation, we applied the RSIL method to measure CO_2_ development from oil mineralization successfully and applied PICRUSt2 for the identification of the potential for hydrocarbon degradation and abundance of two marker genes for activation steps in aerobic hydrocarbon (*alkM* and *ladA*) in our microcosms. By comparison of the actual CO_2_ production trends and rates with the predicted hydrocarbon degradation potential we realized that the predicted potential would be misleading the interpretation and conclusion of our data. We would have missed that the oil degradation potential of the naïve water was vastly overestimated by the functional prediction and that the oil mineralization increased significantly by the application of the seedbank to the anthropogenic oil slick (1:1 oil mixes). This over estimation can be explained by a study that screened 23,446 genomes for the presence of aerobic hydrocarbon-degrading enzymes (19). They detected 2,089 genomes that contained *alkB/M* genes and 2,154 genomes that contained *ladA* genes. However, they reported that only 92 genomes (< 0.5%) of all investigated bacterial genomes had complete or near complete degradation pathways for at least one hydrocarbon compound and that hydrocarbon degradation ability was even less prevalent along archaeal genomes (19). These numbers demonstrate, that predictions on selected genes, although being defined as marker gene, are not proof for hydrocarbon mineralization. Validity and limitations of tools for predicting the metabolic potential of microbial communities based on 16S rRNA gene sequencing data, such as PICRUSt2 (38), FAPOTAX (53), and Tax4Fun (54), are applied in ecological studies but are also highly debated and criticized (55). While these tools might be useful for exploring trends and generating hypotheses, they are not suitable for making definitive functional claims without experimental approval and could even lead to wrong conclusions, like in our case. Main limitations of these tools are the dependence on the reference dataset (56), the assumption that functions are conserved within the same taxa (57), lack of cultured/known taxa and inconsistency with metagenomic data (55). Nonetheless, many studies apply such tools in ecological studies without complementary experiments or measurements, which could lead to wrong conclusions. Thus, studies solely based on such predictive tools should be handled with care.

### Oil from a natural seep serves as a seedbank to enhance hydrocarbon biodegradation

The adapted water from the tar pit contained ∼5 times higher concentrations of dissolved organic carbon than the naïve water, because oil and oil weathering are associated with increased dissolved organic carbon content (58). The dissolved organic carbon content varies largely in rivers worldwide with an average concentration of 10.4 mg L^-1^(59), close to the concentration in the naïve water (14.42 mg L^-1)^. Similar to crude oil, dissolved organic carbon is a very complex organic substrate which cannot be labeled, but its degradation can also be measured with reverse stable isotope labeling (33). In fact, contributions from dissolved organic carbon degradation are also considered in this study, meaning that not necessarily all measured CO_2_ molecules were liberated solely from hydrocarbon mineralization.

It is quite common in oil-contaminated environments that the microbes actively involved in hydrocarbon degradation are not necessarily the most abundant taxa, as we also observed for the microcosm communities, that were different to the initial environmental samples. Some studies have pointed out that the taxonomic abundance alone does not always correlate with functional gene abundance (60, 61). We observed that the microbial community from the tar pit is not exclusively adapted to the degradation of heavy oil, because incubation with light oil resulted in even higher CO_2_ production rates than the heavy oil. This is plausible because light oils generally contain greater proportions of easily biodegradable compounds, such as aliphatic short-chain n-alkanes, while these compounds are mainly depleted in heavy oil due to prior weathering and biodegradation (21). A study on marine bacterial communities incubated four different water samples and one sediment sample with five different types of petroleum (extra light, light, medium, heavy and extra heavy) in batch reactors. Samples were taken from the northwest Gulf of Mexico and authors reported that the sampling sites were not far from oil wells and a natural seepage. Microorganism in all four communities were able to degrade the petroleum types from extra-light to heavy, only the sediment community was capable of degrading the extra-heavy fraction and the abundance of low-abundant taxa increased with oil exposure (62). Another metagenomic study in the Baltic Sea (marine) showed that microbial communities harbored genes for the degradation of long-chain alkanes (C20–C32) and some aromatic compounds (63). Both studies also demonstrated that oil degrading communities can deal with a broader range of different hydrocarbon chemicals.

The result, that light oil was degraded four times faster with aid of microbes from the adapted water and about four times more total CO_2_ was produced compared to naïve water with light oil incubations, showed that the adapted water already contained a very actively degrading microbial community and that it is not only attached to the oil slick but the water itself. This is particular interesting since light oils are associated with higher gasoline and diesel content, which are the most consumed petroleum products worldwide, therefore being major contributors to anthropogenic oil spills. It emphasizes the potential seed bank application, utilizing adapted microbes for freshwater spills. Application of the oil slick from the natural oil seep, as a seed bank, to the light oil slick in naïve water increased the hydrocarbon mineralization rate by 32%. This was in the range of previously found 13-70% increased biodegradation rates after injection of hydrogen peroxide and nutrients after the Exxon Valdez oil tanker spill in the marine Prince William Sound, Alaska. The 70% biodegradation rate occurred only during summer (64), so that temperature effects strongly modulated degradation capacity. The here reported anaerobic CO_2_ production rate from heavy oil in the adapted water (0.002 mM day^-1^) was 2.2 times faster (when interpolated to 0.78 mM year^-1^) than the reported rates of 0.35 mM CO_2_ year^-1^ in a previous published study on anaerobic oil degradation. In this study, samples from an unproduced light oil reservoir and its indigenous microbial communities from the formation water and water droplets entrapped in the oil were studied (30). This was not surprising, as cell numbers and microbial activities are expected to be much lower in a subsurface oil reservoir compared to the quite shallow and open tar pit in Hänigsen.

The use of naïve or adapted water showed no big difference for oil mineralization when heavy oil was added, suggesting that the heavy oil already contained adapted microbes entrapped in water droplets, from the adapted water. It has been reported before that tiny water droplets, of only a few microliters, are entrapped in heavy oil (bitumen) and are containing diverse and actively oil degrading microbial communities (31, 65–67) with proportional large core communities of up to 68.0 ± 19.9% (68). By the water inclusions in the oil slick from the natural oil seep, the oil slick may serve as a protection for the microbiome during the introduction into a new ecosystem or to new environmental conditions, here the naïve water. This protection could support the succession of the seed bank during the transition and initial priming of the oil spill during bioremediation.

A potential bioremediation strategy of anthropogenic oil spills into freshwater could involve leveraging small volumes of collected oil slicks from a natural oil seep together with its seedbank into the freshly contaminated environment. By this introduction, the seep-adapted microorganisms could be protected within their native hydrocarbon “habitat”. This approach may allow these pre-adapted organisms to establish themselves in the new spill environment, thereby priming or accelerating the degradation of the contaminating oil. Such a strategy combines the advantages of using naturally selected, hydrocarbon-degrading microbes with minimal environmental disturbance, potentially enhancing the efficiency and resilience of oil spill remediation efforts. It may seem odd to treat an oil spill with the introduction of even more and weathered heavy oil, but there are advantages: No complicated and expensive pre-cultivations of microbial communities are needed, as simply taken out of the nature. Moreover, the risk that organisms become domesticated, attenuated or alter their composition or behavior during cultivation is reduced. Long-term cultivation of microbial communities for bioremediation of oil-contaminated environments involves several cost components, including infrastructure, labor, maintenance, and quality control (69, 70). Artificial and uncontrolled shifts in long term cultivated microbial communities are well known, and can include the loss of enzymes, or biosynthesis of specific compounds such as amino acids, loss of mismatch repair, more tolerance to changing redox conditions, loss of utilization of certain substrates (71).

## Data archiving statement

All data is provided in the supplement material and raw 16S rRNA gene amplicon sequences were uploaded to the NCBI database under BioProject PRJNA1365740.

## Acknowledgements

The authors want to thank Stephan Lütgert from Deutsches Erdölmuseum Wietze, Thomas Degro, Stefan und Silke Auerbach from the Hänigsen Teerkuhlen Museum for support during sampling and their expertise and passion about crude oil in Germany. We thank Astrid Dannehl for assistance during the sampling trip and Daria Baikova for assistance during dissolved organic carbon determination. The authors wish also to thank Rainer Meckenstock for providing laboratory facilities and equipment. Many thanks to Erling Rykkelid, Joachim Rinna and Aker BP ASA (Norway) for the provision of crude light oil and its chemical composition data. We thank Torsten Schmidt from Instrumental Analytical Chemistry (IAC), University of Duisburg-Essen for access to the TOC-L total organic carbon analyzer.

## References

1. K. A. Kvenvolden, C. K. Cooper, Natural seepage of crude oil into the marine environment. Geo-Marine Letters 23, 140–146 (2003).

2. A. Jernelöv, The Threats from Oil Spills: Now, then, and in the future. Ambio 39, 353–366 (2010).

3. L. Voskuhl, J. Rahlff, Natural and oil surface slicks as microbial habitats in marine systems: A mini review. Front Mar Sci 9, (2022).

4. Y. Dong, Y. Liu, C. Hu, I. R. MacDonald, Y. Lu, Chronic oiling in global oceans. Science 376, 1300–1304 (2022).

5. E. D’Ugo et al., Integration of satellite surveillance and metagenomics for the monitoring and protection of water basins from oil spills. Environmental Advances 15, (2024).

6. S. Rajendran et al., Monitoring oil spill in Norilsk, Russia using satellite data. Sci Rep 11, 3817 (2021).

7. C. Pietrapertosa et al., Hyperspectral images to monitor oil spills in the River Po. B Geofis Teor Appl 57, 31–42 (2016).

8. J. Nriagu, E. A. Udofia, I. Ekong, G. Ebuk, Health Risks Associated with Oil Pollution in the Niger Delta, Nigeria. Int J Environ Res Public Health 13, (2016).

9. A. Jernelov, The threats from oil spills: now, then, and in the future. Ambio 39, 353–366 (2010).

10. R. Dollhopf, M. Durno (2011) Kalamazoo River\Enbridge Pipeline Spill 2010. in International Oil Spill Conference Proceedings.

11. D. Kvočka, D. Žagar, P. Banovec, A Review of River Oil Spill Modeling. Water 13, 1620 (2021).

12. A. Carpenter, Oil pollution in the North Sea: the impact of governance measures on oil pollution over several decades. Hydrobiologia 845, 109–127 (2019).

13. D. P. French-McCay, Development and application of an oil toxicity and exposure model, OilToxEx. Environ. Toxicol. Chem. 21, 2080–2094 (2002).

14. Z. Asif, Z. Chen, C. An, J. Dong, Environmental Impacts and Challenges Associated with Oil Spills on Shorelines. Journal of Marine Science and Engineering 10, 762 (2022).

15. I. B. Ivshina et al., Oil spill problems and sustainable response strategies through new technologies. Environmental Science: Processes & Impacts 17, 1201–1219 (2015).

16. J. R. Bragg, R. C. Prince, E. J. Harner, R. M. Atlas, Effectiveness of bioremediation for the Exxon Valdez oil spill. Nature 368, 413–418 (1994).

17. S. Louca, F. Mazel, M. Doebeli, L. W. Parfrey, A census-based estimate of Earth’s bacterial and archaeal diversity. PLOS Biology 17, e3000106 (2019).

18. V. Khot et al., CANT-HYD: A Curated Database of Phylogeny-Derived Hidden Markov Models for Annotation of Marker Genes Involved in Hydrocarbon Degradation. Front Microbiol 12, 764058 (2021).

19. M. R. Somee et al., Genome-resolved analyses show an extensive diversification in key aerobic hydrocarbon-degrading enzymes across bacteria and archaea. BMC Genomics 23, 690 (2022).

20. J. T. Lennon, S. E. Jones, Microbial seed banks: the ecological and evolutionary implications of dormancy. Nat Rev Microbiol 9, 119–130 (2011).

21. I. M. Head, D. M. Jones, W. F. M. Röling, Marine microorganisms make a meal of oil. Nat Rev Microbiol 4, 173–182 (2006).

22. W. F. M. Röling et al., Robust Hydrocarbon Degradation and Dynamics of Bacterial Communities during Nutrient-Enhanced Oil Spill Bioremediation. Applied and Environmental Microbiology 68, 5537–5548 (2002).

23. H. P. Bacosa et al., Polycyclic aromatic hydrocarbons (PAHs) and putative PAH-degrading bacteria in Galveston Bay, TX (USA), following Hurricane Harvey (2017). Environmental Science and Pollution Research 27, 34987–34999 (2020).

24. A. K. Williams, H. P. Bacosa, A. Quigg, The impact of dissolved inorganic nitrogen and phosphorous on responses of microbial plankton to the Texas City “Y” oil spill in Galveston Bay, Texas (USA). Marine Pollution Bulletin 121, 32–44 (2017).

25. M. Tyagi, M. M. da Fonseca, C. C. de Carvalho, Bioaugmentation and biostimulation strategies to improve the effectiveness of bioremediation processes. Biodegradation 22, 231–241 (2011).

26. C. Gertler, G. Gerdts, K. N. Timmis, P. N. Golyshin, Microbial consortia in mesocosm bioremediation trial using oil sorbents, slow-release fertilizer and bioaugmentation. FEMS Microbiol Ecol 69, 288–300 (2009).

27. Heimatbund Hänigsen im Heimatbund Niedersachsen e.V. (Die Hänigser Teerkuhlen.

28. S. Beilig et al., Coexistence of methanogenesis and sulfate reduction in a sulfate-adapted enrichment culture from an oil reser Methanogenesis and Sulfate Reduction in a Sulfate-Adapted Enrichment Culture From an Oil Reservoir. Applied and Environmental Microbiology AEM00141–25, (2025).

29. P. Mandal, M. Sasaki, “Total acid number reduction of naphthenic acids using supercritical fluid and ionic liquids” in Recent insights in petroleum science and engineering, M. Zoveidavianpoor, Ed. (IntechOpen, London, 2018), 10.5772/intechopen.71812.

30. S. Beilig, M. Pannekens, L. Voskuhl, R. U. Meckenstock, Assessing anaerobic microbial degradation rates of crude light oil with reverse stable isotope labelling and community analysis. Frontiers in Microbiomes 3, (2024).

31. M. Pannekens et al., Microbial degradation rates of natural bitumen. Environ Sci Technol 55, 8700–8708 (2021).

32. T. B. Coplen, Guidelines and recommended terms for expression of stable-isotope-ratio and gas-ratio measurement results. Rapid Commun Mass Spectrom 25, 2538–2560 (2011).

33. X. Dong et al., Quantification of microbial degradation activities in biological activated carbon filters by reverse stable isotope labelling. AMB Express 9, 109 (2019).

34. A. Klindworth et al., Evaluation of general 16S ribosomal RNA gene PCR primers for classical and next-generation sequencing-based diversity studies. Nucleic Acids Res 41, e1 (2013).

35. B. J. Callahan et al., DADA2: High-resolution sample inference from Illumina amplicon data. Nat Methods 13, 581–583 (2016).

36. P. J. McMurdie, S. Holmes, phyloseq: an R package for reproducible interactive analysis and graphics of microbiome census data. PLoS One 8, e61217 (2013).

37. C. Quast et al., The SILVA ribosomal RNA gene database project: improved data processing and web-based tools. Nucleic Acids Res 41, D590–596 (2013).

38. G. M. Douglas et al., PICRUSt2 for prediction of metagenome functions. Nature Biotechnology 38, 685–688 (2020).

39. Clarivate (2025) Web of Science.

40. K. K. Sjøholm et al., Linking biodegradation kinetics, microbial composition and test temperature – Testing 40 petroleum hydrocarbons using inocula collected in winter and summer. Environmental Science: Processes & Impacts 24, 152–160 (2022).

41. V. H. T. Pham, S.-W. Jeong, J. Kim, *Aquabacterium olei* sp. nov., an oil-degrading bacterium isolated from oil-contaminated soil. International Journal of Systematic and Evolutionary Microbiology 65, 3597–3602 (2015).

42. L. Ren et al., Parablastomonas arctica gen. nov., sp. nov., isolated from high Arctic glacial till. International Journal of Systematic and Evolutionary Microbiology 65, 260–266 (2015).

43. M. O. Mafiana, X. H. Kang, Y. Leng, L. F. He, S. W. Li, Petroleum contamination significantly changes soil microbial communities in three oilfield locations in Delta State, Nigeria. Environ Sci Pollut Res Int 28, 31447–31461 (2021).

44. A. Samad et al., Microbial community structural and functional differentiation in capped thickened oil sands tailings planted with native boreal species. Front Microbiol 14, 1168653 (2023).

45. U. Passow, K. Ziervogel, V. Asper, A. Diercks, Marine snow formation in the aftermath of the Deepwater Horizon oil spill in the Gulf of Mexico. Environmental Research Letters 7, (2012).

46. K. Ziervogel, N. D’Souza, J. Sweet, B. Yan, U. Passow, Natural oil slicks fuel surface water microbial activities in the northern Gulf of Mexico. Front Microbiol 5, 188 (2014).

47. H. P. Bacosa et al., Extracellular polymeric substances (EPS) producing and oil degrading bacteria isolated from the northern Gulf of Mexico. PLOS ONE 13, e0208406 (2018).

48. L. Sun et al., The effects of sunlight on the composition of exopolymeric substances and subsequent aggregate formation during oil spills. Marine Chemistry 203, 49–54 (2018).

49. H. P. Bacosa et al., From Surface Water to the Deep Sea: A Review on Factors Affecting the Biodegradation of Spilled Oil in Marine Environment. Journal of Marine Science and Engineering 10, 426 (2022).

50. B. H. Gregson et al., Marine Oil Snow, a Microbial Perspective. Frontiers in Marine Science Volume 8- 2021, (2021).

51. R. Takeshita et al., A review of the toxicology of oil in vertebrates: what we have learned following the Deepwater Horizon oil spill. Journal of Toxicology and Environmental Health, Part B 24, 355–394 (2021).

52. R. Almeda et al., Effects of crude oil exposure on bioaccumulation of polycyclic aromatic hydrocarbons and survival of adult and larval stages of gelatinous zooplankton. PLoS One 8, e74476 (2013).

53. A. Ahmad, H. Fröhlich, Towards clinically more relevant dissection of patient heterogeneity via survival-based Bayesian clustering. Bioinformatics 33, 3558–3566 (2017).

54. F. Wemheuer et al., Tax4Fun2: prediction of habitat-specific functional profiles and functional redundancy based on 16S rRNA gene sequences. Environmental Microbiome 15, 11 (2020).

55. M. S. Matchado et al., On the limits of 16S rRNA gene-based metagenome prediction and functional profiling. Microb Genom 10, (2024).

56. R. J. Wright, M. G. I. Langille, PICRUSt2-SC: an update to the reference database used for functional prediction within PICRUSt2. Bioinformatics 41, (2025).

57. S. Sun, R. B. Jones, A. A. Fodor, Inference-based accuracy of metagenome prediction tools varies across sample types and functional categories. Microbiome 8, 46 (2020).

58. T. S. Catalá, S. Shorte, T. Dittmar, Marine dissolved organic matter: a vast and unexplored molecular space. Applied Microbiology and Biotechnology 105, 7225–7239 (2021).

59. F. Liu, D. Wang, Dissolved organic carbon concentration and biodegradability across the global rivers: A meta-analysis. Science of The Total Environment 818, 151828 (2022).

60. N. Das, B. Bhuyan, P. Pandey, Correlation of soil microbiome with crude oil contamination drives detection of hydrocarbon degrading genes which are independent to quantity and type of contaminants. Environ. Res. 215, 114185 (2022).

61. T. H. Bell et al., A Diverse Soil Microbiome Degrades More Crude Oil than Specialized Bacterial Assemblages Obtained in Culture. Appl Environ Microbiol 82, 5530–5541 (2016).

62. D. Cerqueda-García, J. Q. García-Maldonado, L. Aguirre-Macedo, U. García-Cruz, A succession of marine bacterial communities in batch reactor experiments during the degradation of five different petroleum types. Marine Pollution Bulletin 150, 110775 (2020).

63. J. M. Serrana, B. Dessirier, F. J. A. Nascimento, E. Broman, M. Posselt, Microbial hydrocarbon degradation potential of the Baltic Sea ecosystem. Microbiome 13, 204 (2025).

64. M. C. Boufadel, X. Geng, J. Short, Bioremediation of the Exxon Valdez oil in Prince William Sound beaches. Marine Pollution Bulletin 113, 156–164 (2016).

65. R. U. Meckenstock et al., Oil biodegradation. Water droplets in oil are microhabitats for microbial life. Science 345, 673–676 (2014).

66. L. Voskuhl et al., Indigenous microbial communities in heavy oil show a threshold response to salinity. FEMS Microbiol Ecol 97, (2021).

67. M. Pannekens et al., Densely populated water droplets in heavy-oil seeps. Appl Environ Microbiol 86, (2020).

68. V. S. Brauer et al., Imprints of ecological processes in the taxonomic core community: an analysis of naturally replicated microbial communities enclosed in oil. FEMS Microbiol Ecol 100, (2024).

69. R. Orellana et al., Economic Evaluation of Bioremediation of Hydrocarbon-Contaminated Urban Soils in Chile. Sustainability 14, 11854 (2022).

70. M. I. Matilda, H. S. Samuel, Bioremediation of oil spill: concept, methods and applications. Discover Chemistry 1, 42 (2024).

71. J. Steensels, B. Gallone, K. Voordeckers, K. J. Verstrepen, Domestication of Industrial Microbes. Current Biology 29, R381–R393 (2019).

