## Supplementary text for "A microbial seedbank from a natural oil seep accelerates hydrocarbon degradation in freshwater oil spills"

**Additional method information**

**Cell counting via NovoCyte, Agilent flow cytometer** (Santa Clara, USA)

Samples taken for cell counting were kept on ice during the sampling and were frozen at ‑70 °C until use. The measurement of cell numbers using flow cytometry was not reliably feasible as too many microbial cells stayed attached to the oil and later oil snow and were excluded after filtration (pore size 10 µm) during the sample preparation for the flow cytometry measurement.

**Literature search**

A literature survey was conducted using the Web of Science Core Collection (39) considering publications from 1900 to 2025 to quantify research on oil slicks and their microbial aspects in both marine and freshwater systems. Four main categories were defined for targeted searches: 1) general studies on oil slicks in marine environments, 2) studies in marine environments emphasizing microbial ecology, hydrocarbon degradation, or bioremediation, 3) general studies on oil slicks in lakes, rivers, streams, or ponds, 4) studies in freshwater systems addressing microbial responses, biodegradation, or bioremediation. Advanced query searches were performed using the Topic Search (TS) field in Web of Science. Boolean operators were applied to combine oil-related terms with environmental and microbial terms. For each query, the following metadata were retrieved: total publications, review articles, early access articles, open access articles, and enriched cited references. The queries for each category were as follows:

1) Marine slicks: TS=((""oil slick*"" OR ""oil spill*"" OR ""oil film*"" OR ""hydrocarbon contamination"") AND (marine OR ocean* OR sea OR estuar*))
2) Marine oil slicks with microbial focus: TS=((""oil slick*"" OR ""oil spill*"" OR ""oil film*"" OR ""hydrocarbon contamination"") AND (marine OR ocean* OR sea OR estuar*) AND (microb* OR bacteri* OR degrad* OR bioremediat* OR ""microbial ecology""))
3) Freshwater oil slicks: TS=((""oil slick*"" OR ""oil spill*"" OR ""oil film*"" OR ""hydrocarbon contamination"") AND (freshwater OR lake* OR river* OR stream* OR pond*))
4) Freshwater oil slicks with microbial focus:	TS=((""oil slick*"" OR ""oil spill*"" OR ""oil film*"" OR ""hydrocarbon contamination"") AND (freshwater OR lake* OR river* OR stream* OR pond*) AND (microb* OR bacteri* OR degrad* OR bioremediat* OR ""microbial ecology""))

**Mass balance for assessment of O_2_ consumption and to calculate how much oil was degraded**

Following assumptions were applied for O_2_ consumption and oil mass degradation calculations. CH_2_ was considered as the smallest building block of oil (with the oxidation state of -2), the following redox equation was formulated for the aerobic mineralization of oil:

$$unbalanced equation: CH_{2}+O_{2}\to CO_{2}+H_{2}O$$

$$Oxidation: CH_{2}+ 2 H_{2}O\to CO_{2}+6e^{-}+6H^{+} |\cdot2$$

$$Reduction: O_{2}+4e^{-}+4H^{+}\to2H_{2}O |\cdot3$$

$$Ox+Red: 2 CH_{2}+3 O_{2}\to2 CO_{2}+2H_{2}O$$

Oil is mineralized to CO_2_ in a ratio of 1:1, which means that 1 mol oil is converted to 1 mol CO_2_. The total volume of water and oil in the microcosm was approximately 100 mL, therefore:

$12 \frac{\mathrm{mmol}}{L}\cdot0.1 L=1.2 mmol CO_{2} \mathrm{produced}$

To calculate the mass of degraded oil from mmol to gram, the following equation with M(CH_2_) = 14 g mol^-1^ was applied:

$n=\frac{m}{M} \leftrightarrow m=n\cdot M$

$m=1.2\cdot{10}^{-3} mol\cdot14 \frac{g}{\mathrm{mol}}=16.8\cdot{10}^{-3}g$

$m=16.8 mg$

Next, the amount of consumed oxygen for oil degradation was estimated. First, the volume of the headspace inside the microcosm was determined by filling up a 250 mL Schott flask to the rim with water. A butyl stopper was inserted, displacing small amounts of water. The butyl stopper was removed and the water volume was measured with a measuring cylinder. To obtain the headspace volume in each microcosm, 100 mL was deducted to account for the water and oil portion inside the microcosms. The determined headspace volume was 210 mL. Oxygen makes up about 20 % of atmospheric air, therefore 42 mL O_2_ were present in the headspace of the microcosms and available for oil degradation. From the balanced redox equation, it can be seen that for the mineralization of 1 mol oil, 3/2 mol oxygen are needed. Therefore, the used oxygen equates to:

$n\left( O_{2} \right)=\frac{3}{2}n\left( \mathrm{oil} \right)=\frac{3}{2}\cdot1.2\cdot{10}^{-3} mol=1.8 mmol O_{2}$

To convert 1.8 mmol oxygen to mL, the following equation was used with V_m_ = 22.4 L mol^-1^ (V_m_ is the molar volume and can be obtained from the ideal gas equation $pV=nRT \leftrightarrow\frac{V}{n}=\frac{\mathrm{RT}}{p}$):

$n\left( O_{2} \right)=\frac{V\left( O_{2} \right)}{V_{m}} \leftrightarrow V\left( O_{2} \right)=n\left( O_{2} \right)\cdot V_{m}$

$V\left( O_{2} \right)=1.8 \cdot{10}^{-3} mol\cdot22.4 \frac{L}{\mathrm{mol}}$

$V\left( O_{2} \right)=0.04032 L=40.32 mL$

Next, the percentile of remaining oxygen in the headspace was calculated as followed:

$\mathrm{remaining}O_{2} in headspace in \%=\left( 1-\frac{40.32 mL}{42 mL} \right)\cdot100\%=4 \%$

The required amount of oxygen for the degradation of 1 g oil was calculated from the relation of used oxygen (in mL) to oil (in mg) according to:

$\frac{40.32 mL {(O}_{2})}{16.8 mg (oil)}=2.4\frac{\mathrm{mL}{(O}_{2})}{mg (oil)} =2.4\frac{L (O_{2})}{g (oil)}$

This shows that the complete aerobic degradation of oil was not possible in such small microcosms due to electron acceptor limitations.

**16S rRNA gene amplicon results of the positive and negative controls**

Mock ZymoBIOMICS™ Microbial Community Standard (Catalog No. D6300) was applied as positive control to validate DNA extraction methods, Library preparation and Illumina MiSeq sequencing analyzed with DADA2 pipeline. The measured relative abundances of the mock community are close but not exact to the given theoretical relative abundance as expected (Table S1 and Figure S1). *Listeria* (ASV10) and *Staphylococcus* (ASV08), *Salmonella* (ASV13) *Escherichia-Shigella* (ASV14) and *Enterococcus* (ASV19) were about 65%, 32%, 25% 23% and 22% over represented, respectively. While *Bacillus* (ASV12), *Pseudomonas* (ASV40) and Limosilactobacillus (ASV23 and 62) were underrepresented by 28%, 20% and 82% respectively. A reason for this might be that the DNA was extracted in unequal proportions or a primer bias, meaning that employed primers bind more likely to e.g. Staphylococcus and less likely to Pseudomonas. The remaining missing 0.66% DNA in the measured mock community could not be assigned to ASVs from the samples and are rated as cross contaminations with low reads.

Negative controls were proceeded along the whole workflow in order to test the reagents purities and clean working-style. A clean working-style was confirmed by read numbers below 100 looking very similar to the positive control. The top 250 organisms of the negative control accounted for 91.6 % of total abundance and could be mainly assigned to the positive control (Figure S1).

**Table S1** Theoretically expected composition of the Mock community applied as positive control (left) and the true outcome from workflow (right) after Mock was used for DNA extraction, Library preparation and Illumina MiSeq sequencing analyzed with DADA2 pipeline. The Deviation in % is displayed in the last column, values above indicate a over estimation by the method and values below 100% an underestimation of the organism.

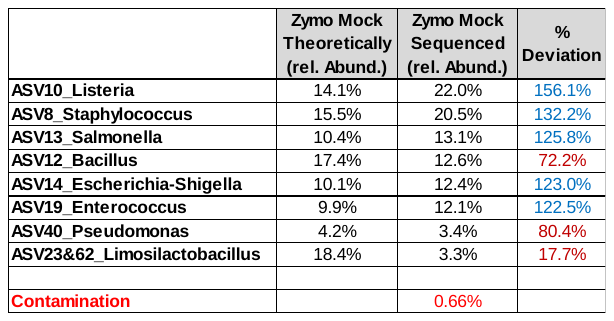

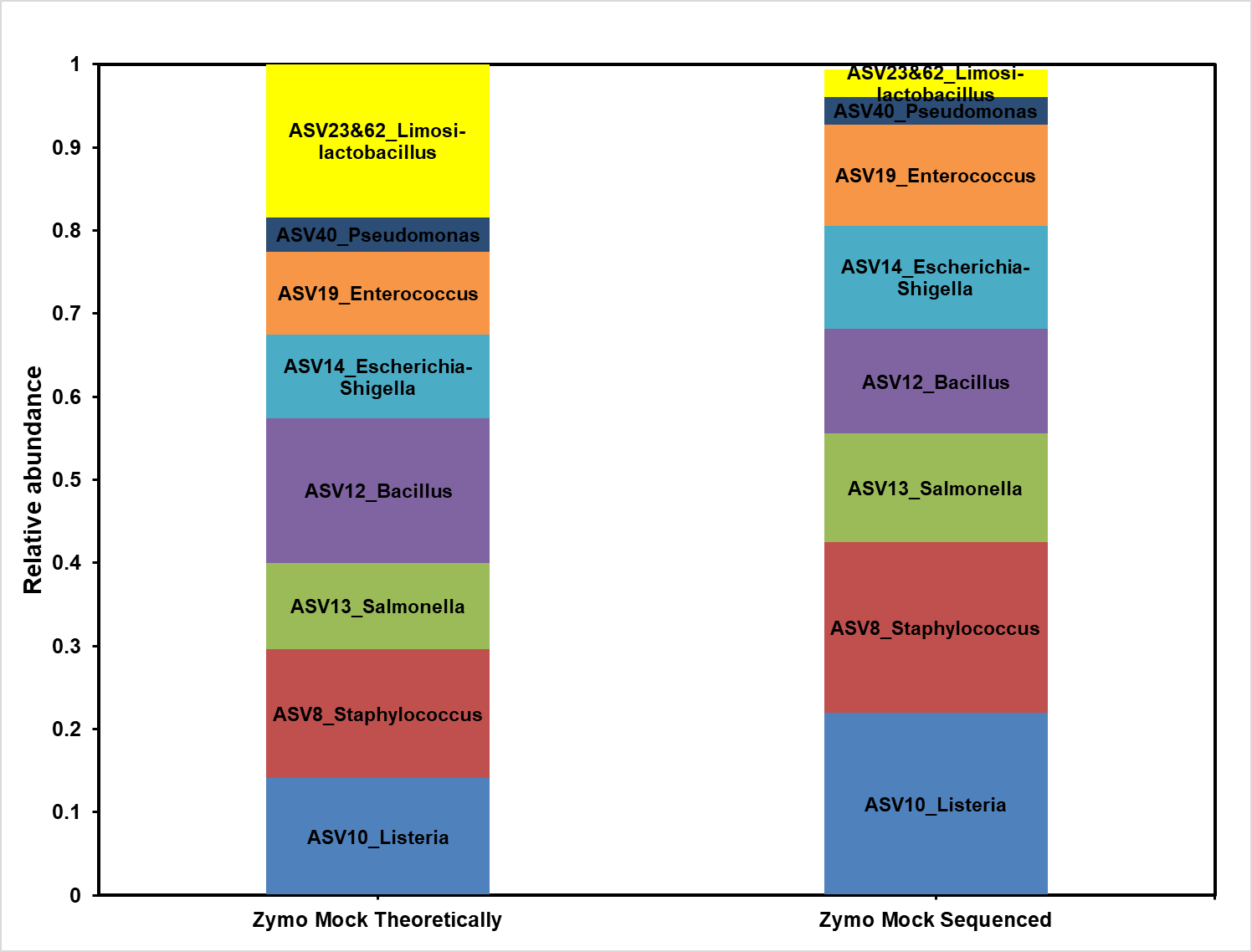

Figure S1 Bar plots of the theoretic expected composition of the Mock community applied as positive control (left) and the true outcome from workflow (right) after Mock was used for DNA extraction, Library preparation and Illumina MiSeq sequencing analyzed with DADA2 pipeline.

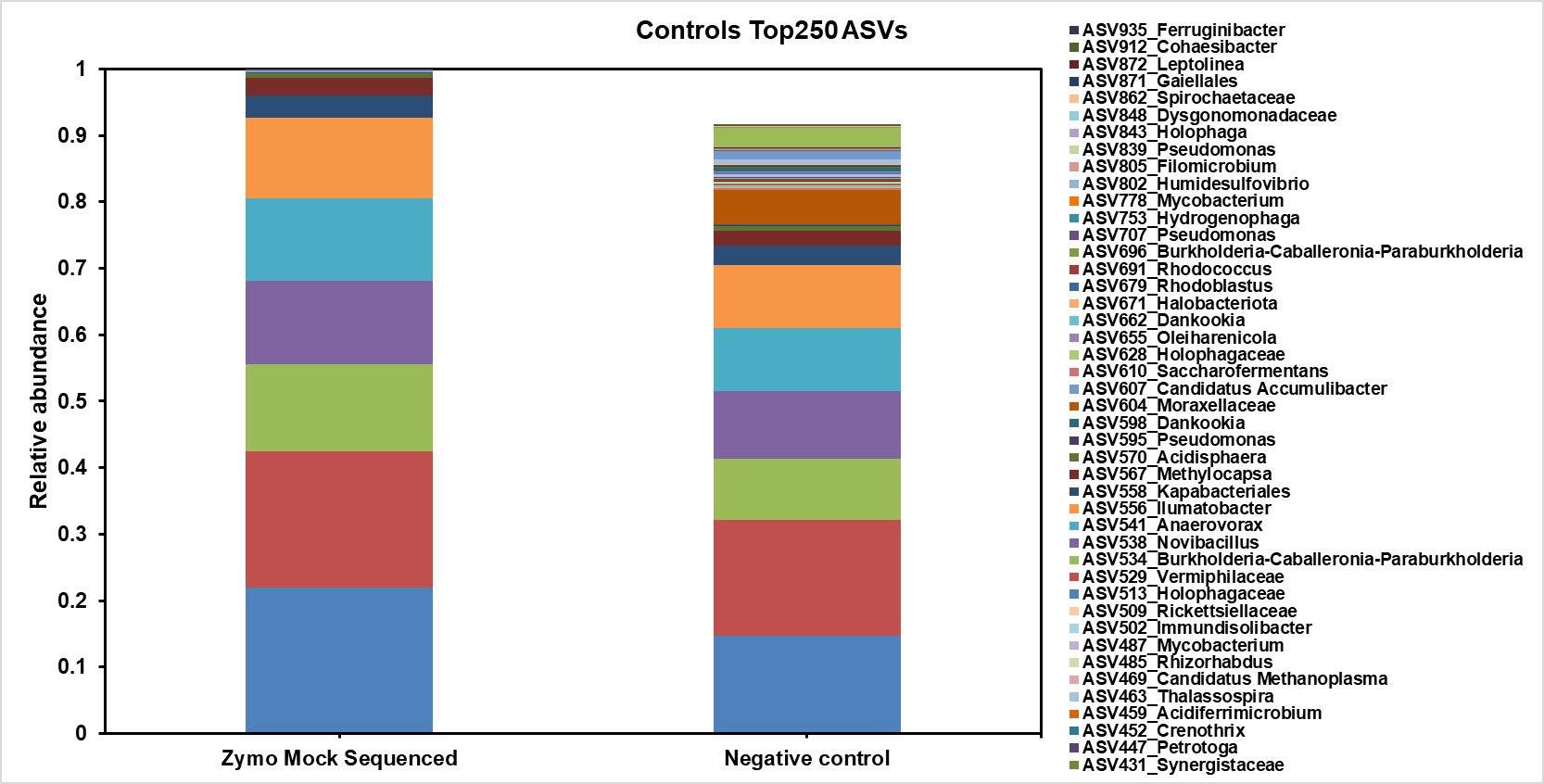

Figure S2 Bar plot of the negative control (right) compared to the positive control.

Bray-Curtis dissimilarity plot for the controls and microbial communities from each test condition demonstrated that all controls (top left corner green triangles) looked different to the samples.

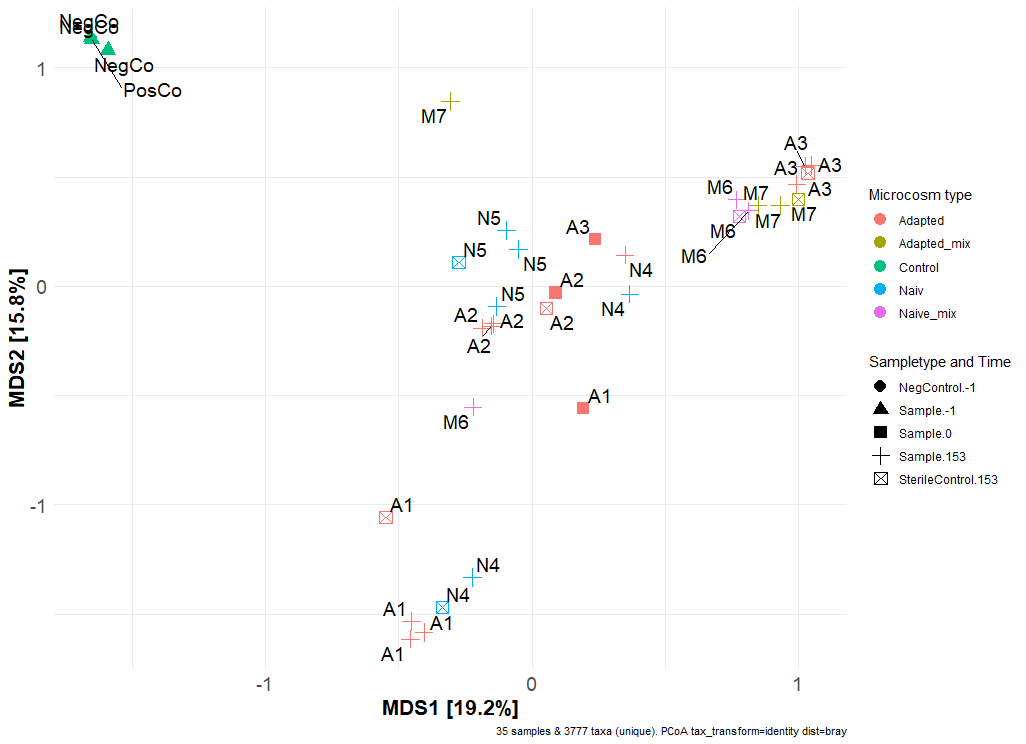

Figure S3 Compositional community analysis based on Bray-Curtis dissimilarity. MDS plot for the controls and microbial communities from each test condition. This plot shows that the negative controls (green) contain very different microbial communities to the samples and that they completely overlap with the positive control.

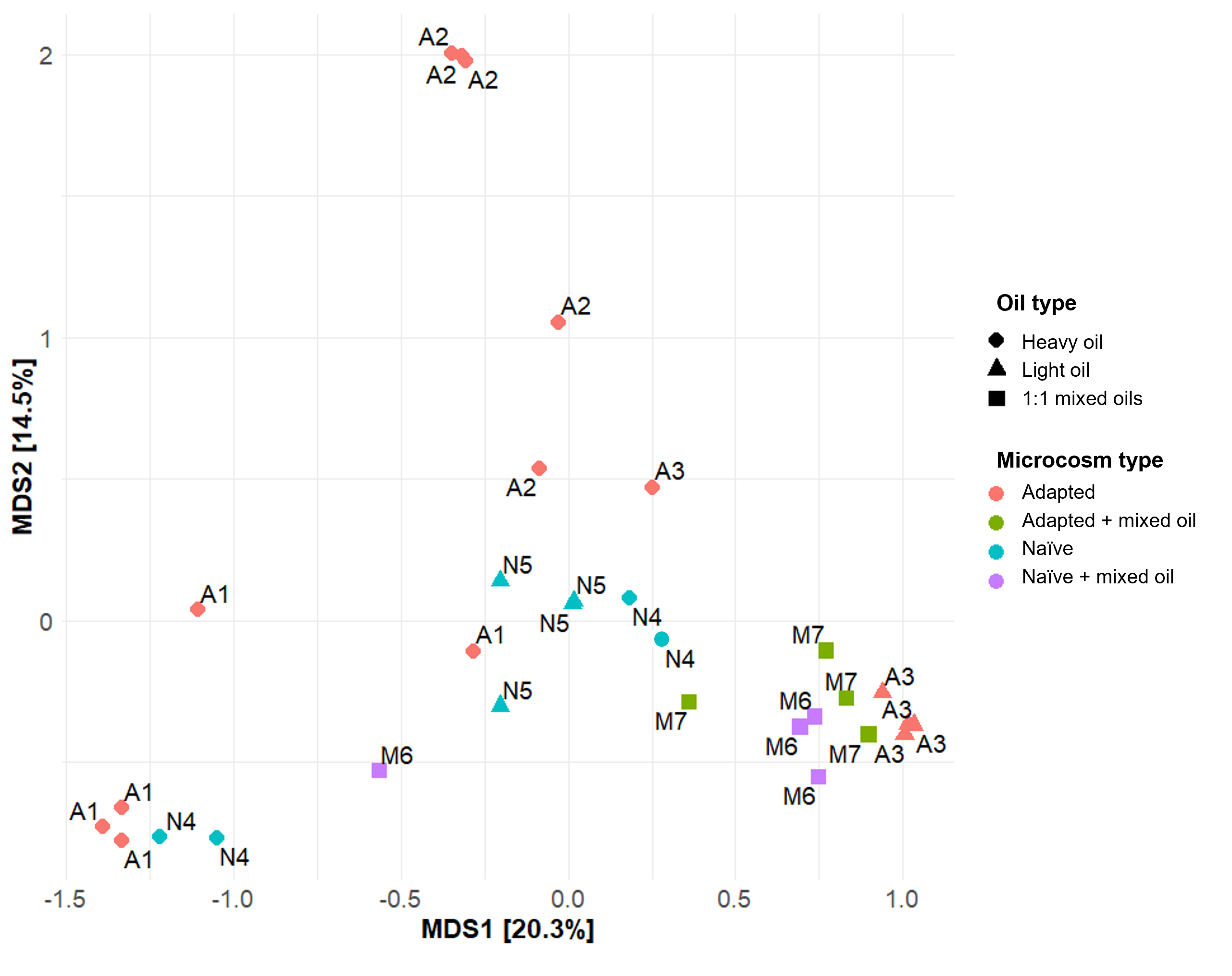

Figure S4 Compositional community analysis based on Bray-Curtis dissimilarity. MDS plot for the shapes indicate the type of applied oil and the colors represent the microcosm type. This plot shows that the mixed oil treatments in adapted (M7) and in naïve (M6) water were most similar to the adapted water treated with light oil, which also showed highest mineralization rates.

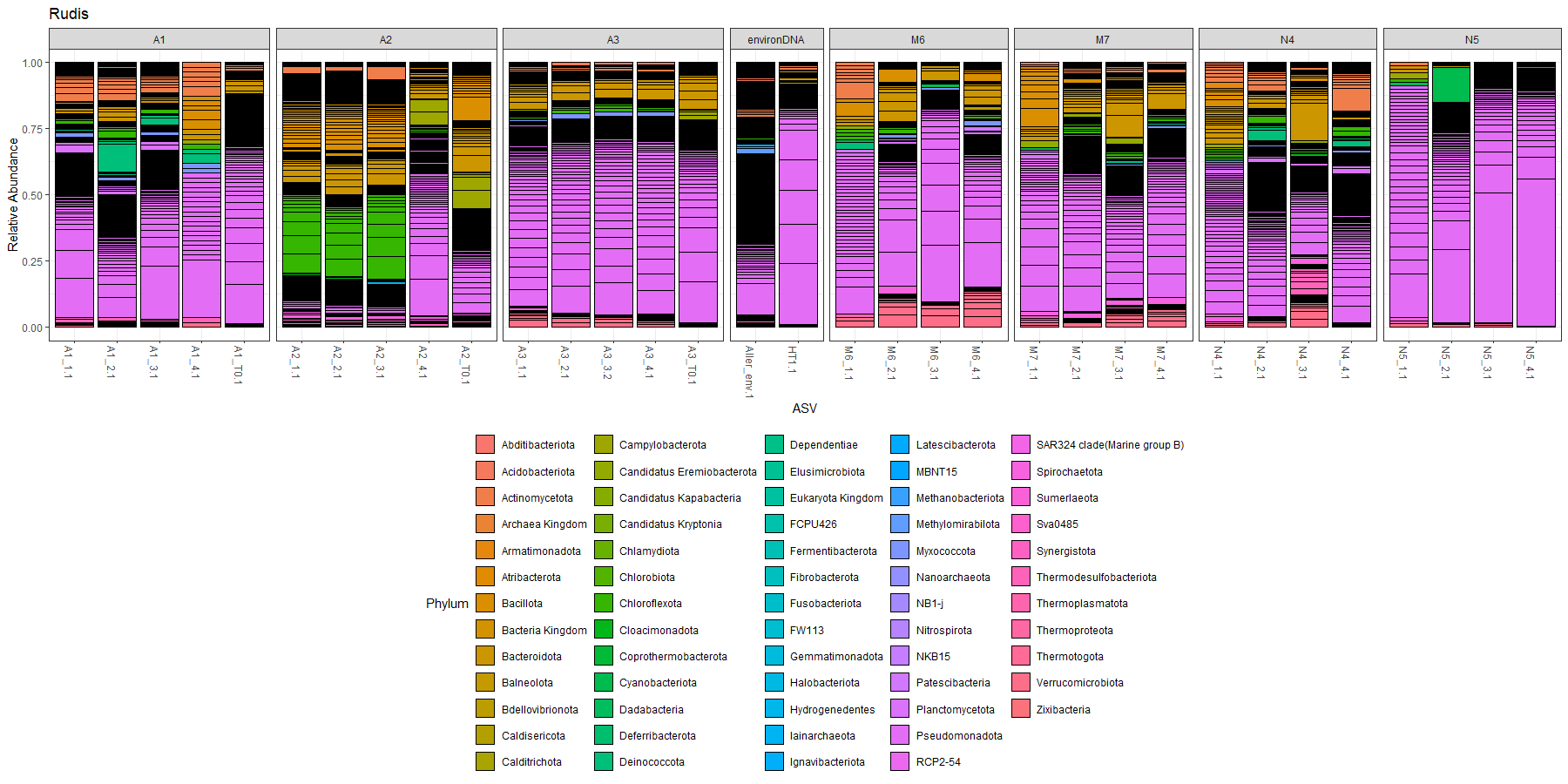

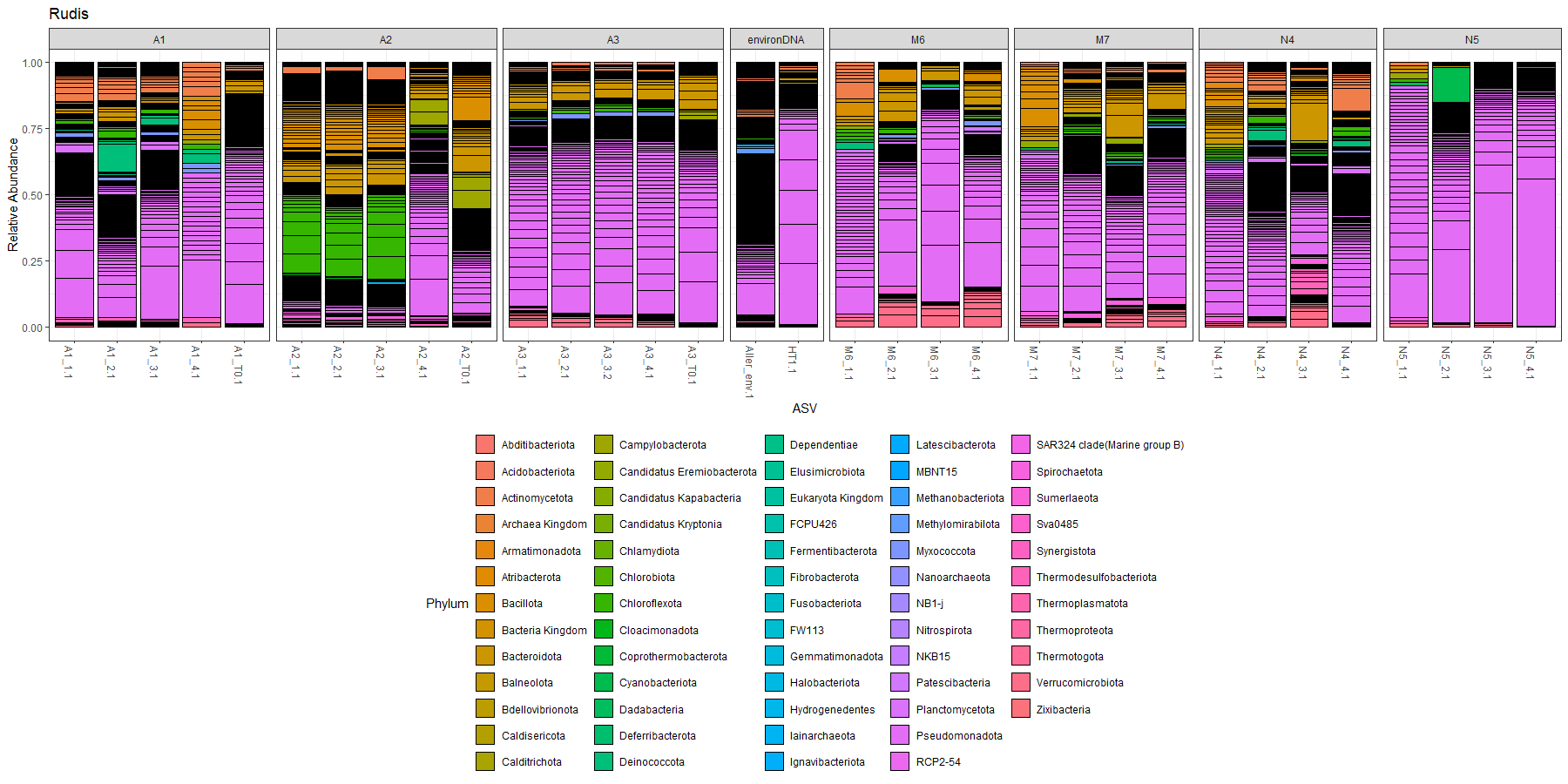

Figure S5 16S rRNA gene amplicon composition of the individual samples of the biological replicates, and the autoclaved controls (*_4.1).

Different to potential genes for oil degradation (see Table S2), potential genes for central metabolism (Table S3 and Figure S6) where quite homogeneously distributed along all samples, identified with PICRUSt2. Approximately, 6% of all detected pathways could be assigned to central metabolisms. Following 11 genes chosen that are involved in hydrocarbon degradation:

1.	aerobic activation steps: alkane 1-monooxygenase (alkB1_2, alkM)

2.	aerobic activation steps: long-chain alkane monooxygenase (ladA)

3.	Aromatic hydrocarbons (anaerobic pathways): benzoate-CoA ligase (badA)

4.	benzene/toluene/chlorobenzene dioxygenase subunit alpha (todC1, bedC1, tcbAa)

5.	phenol/toluene 2-monooxygenase (NADH) P0/A0 (dmpK, poxA, tomA0)

6.	benzoate/toluate 1,2-dioxygenase subunit alpha (benA-xylX)

7.	dihydroxycyclohexadiene carboxylate dehydrogenase (benD-xylL)

8.	catechol 1,2-dioxygenase (catA)

9.	muconate cycloisomerase (catB)

10.	catechol 2,3-dioxygenase (dmpB, xylE)

11.	cyclohexa-1,5-dienecarbonyl-CoA hydratase (dch)

Table S2 Pathway names and explanations for potential oil degradation that were selected and detected via PICRUSt2.

| **Function** | **Pathway** |
| --- | --- |
| degradation of 1,2-Dichlorethane | 12DICHLORETHDEG-PWY |
| degradation of 1,4-Dichlorbenzole | 14DICHLORBENZDEG-PWY |
| degradation of 3-Hydroxyphenylacetate | 3-HYDROXYPHENYLACETATE-DEGRADATION-PWY |
| degradation of 4-Hydroxymandelate | 4-HYDROXYMANDELATE-DEGRADATION-PWY |
| Ortho-cleavage of Catechol | CATECHOL-ORTHO-CLEAVAGE-PWY |
| Gallat-degradation | GALLATE-DEGRADATION-I-PWY |
| Gallat-degradation | GALLATE-DEGRADATION-II-PWY |
| degradation of 4-Hydroxycoumarine-like compounds | HCAMHPDEG-PWY |
| degradation of m-Kresole | M-CRESOL-DEGRADATION-PWY |
| degradation of Methygallate | METHYLGALLATE-DEGRADATION-PWY |
| degradation of Pentachlorphenole (PCP) | PCPDEG-PWY |
| Ortho-cleavage of Protocatechuate | PROTOCATECHUATE-ORTHO-CLEAVAGE-PWY |
| Toluol-degradation via 3-Hydroxy pathway | TOLUENE-DEG-3-OH-PWY |

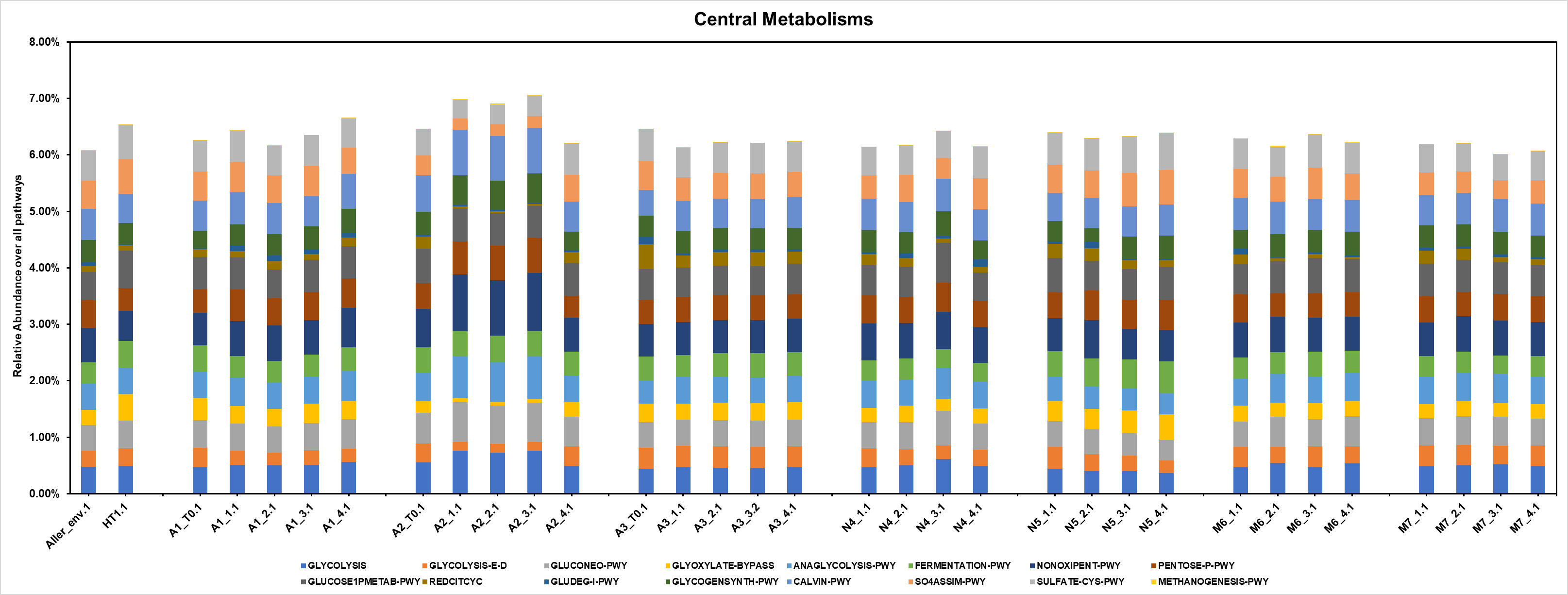

Figure S6 Distribution of the potential fraction of pathways that are involved in the central energy metabolism. These are evenly distributed along all investigated samples.

Table S3 Pathway names and explanations for potential centra metabolisms that were selected and detected via PICRUSt2.

| **Name** | **Function** | **Pathway** |
| --- | --- | --- |
| Glycolysis: Embden-Meyerhof-Parnas (EMP) pathway | Breaks down glucose → pyruvate, generating ATP and NADH | GLYCOLYSIS |
| Entner-Doudoroff pathway | Alternative glycolytic pathway → glucose → pyruvate + glyceraldehyde-3-phosphate | GLYCOLYSIS-E-D |
| Gluconeogenesis pathway | Synthesizes glucose from non-sugar precursors like pyruvate, lactate, amino acids | GLUCONEO-PWY |
| Glyoxylate-Bypass | Modified TCA cycle used when growing on 2-carbon compounds (e.g. acetate) | GLYOXYLATE-BYPASS |
| Anaerobic glycolysis | ATP via substrate-level phosphorylation without oxygen | ANAGLYCOLYSIS-PWY |
| Fermentation | Converts pyruvate into organic end products (e.g. lactate, ethanol, acetate) to regenerate NAD⁺ | FERMENTATION-PWY |
| Non-oxidative pentose phosphate pathway | Interconversion of sugar phosphates (e.g. xylulose-5-phosphate ↔ fructose-6-phosphate) | NONOXIPENT-PWY |
| Pentose phosphate pathway (oxidative branch) | Converts glucose-6-phosphate → ribose-5-phosphate + NADPH | PENTOSE-P-PWY |
|  | Metabolism of glucose-1-phosphate, a key intermediate in: Glycogen synthesis/degradation Conversion to glucose-6-phosphate → enters glycolysis or pentose phosphate pathway | GLUCOSE1PMETAB-PWY |
| Reductive citric acid cycle (reverse TCA) | Fixes CO₂ into organic carbon (opposite of oxidative TCA) | REDCITCYC |
| Glutamate degradation | Links amino acid catabolism with central carbon metabolism (TCA cycle) | GLUDEG-I-PWY |
| Glycogen synthesis | Synthesis of glycogen, a storage form of glucose | GLYCOGENSYNTH-PWY |
| Calvin-Benson-Bassham cycle | Carbon fixation: CO₂ → sugars | CALVIN-PWY |
| Sulfate assimilation | Supplies reduced sulfur for amino acid and cofactor biosynthesis | SO4ASSIM-PWY |
| Incorporates sulfide into cysteine | Key pathway for sulfur-containing amino acid biosynthesis | SULFATE-CYS-PWY |
| Methane production | Terminal step in anaerobic carbon cycle. | METHANOGENESIS-PWY |
